# Assessment of Emerging Pretraining Strategies in Interpretable Multimodal Deep Learning for Cancer Prognostication

**DOI:** 10.1101/2022.11.21.517440

**Authors:** Zarif L. Azher, Anish Suvarna, Ji-Qing Chen, Ze Zhang, Brock C. Christensen, Lucas A. Salas, Louis J. Vaickus, Joshua J. Levy

**Affiliations:** Thomas Jefferson High School for Science and Technology, Alexandria, VA; Cancer Biology Graduate Program, Dartmouth College Geisel School of Medicine, Hanover, NH; Program in Quantitative Biomedical Sciences, Dartmouth College Geisel School of Medicine, Hanover, NH; Department of Epidemiology, Dartmouth College Geisel School of Medicine, Hanover, NH; Department of Molecular and Systems Biology, Dartmouth College Geisel School of Medicine, Hanover, NH; Department of Community and Family Medicine, Dartmouth College Geisel School of Medicine, Hanover, NH; Integrative Neuroscience at Dartmouth (IND) graduate program, Dartmouth College Geisel School of Medicine, Hanover, NH; Emerging Diagnostic and Investigative Technologies, Department of Pathology and Laboratory Medicine, Dartmouth Health, Lebanon, NH; Department of Dermatology, Dartmouth Health, Lebanon, NH

**Keywords:** multimodal, survival, DNA methylation, whole slide images, gene expression, machine learning, graph neural networks, tumor infiltrating lymphocytes

## Abstract

Deep learning models have demonstrated the remarkable ability to infer cancer patient prognosis from molecular and anatomic pathology information. Studies in recent years have demonstrated that leveraging information from complementary multimodal data can improve prognostication, further illustrating the potential utility of such methods. Model interpretation is crucial for facilitating the clinical adoption of deep learning methods by fostering practitioner understanding and trust in the technology. However, while prior works have presented novel multimodal neural network architectures as means to improve prognostication performance, these approaches: 1) do not comprehensively leverage biological and histomorphological relationships and 2) make use of emerging strategies to “pretrain” models (i.e., train models on a slightly orthogonal dataset/modeling objective) which may aid prognostication by reducing the amount of information required for achieving optimal performance. Here, we develop an interpretable multimodal modeling framework that combines DNA methylation, gene expression, and histopathology (i.e., tissue slides) data, and we compare the performances of crossmodal pretraining, contrastive learning, and transfer learning versus the standard procedure in this context. Our models outperform the existing state-of-the-art method (average 11.54% C-index increase), and baseline clinically driven models. Our results demonstrate that the selection of pretraining strategies is crucial for obtaining highly accurate prognostication models, even more so than devising an innovative model architecture, and further emphasize the all-important role of the tumor microenvironment on disease progression.

## Background

### Cancer Prognostication and Machine Learning

Despite improvements in disease management options and global public health efforts, nearly half of Americans will develop cancer in their lifetime and cancer is the second leading cause of death worldwide. In 2022 alone, it is estimated that over 600,000 individuals in the United States will die from cancer [1]. As each individual struggles with the disease differently, cancer care has become increasingly personalized. As such, researchers and clinicians are working collaboratively to develop and apply treatments tailored to the specific disease progression of individual patients. The patient’s prognosis indicates the eventual disease outcome or risk of death, which is paramount for determining effective therapies [2]. Prognostication is a difficult and often subjective task which relies on information such as disease stage and molecular markers (ex; mutation status), where practitioners must rely heavily upon a gestalt impression of the patient’s risk profile for clinical decision-making. Inferring this information can be especially challenging when considering the troves of biomedical information that are now available as enabled through the transformative impact of big data and subsequent high throughput processing methods. Thus, there is a lack of prognostically relevant clinically-adopted molecular information derived from these data. Machine learning (ML) is a discipline where models are optimized by task-specific objective functions to make sense of large datasets. Deep learning models are particularly effective and leverage artificial neural networks (ANNs) to make predictions based on input data [3]. As data availability has proliferated through large-scale consortiums such as The Cancer Genome Atlas (TCGA), ML techniques have been applied to integrate a wide range of input data modalities for prognostication [4], such as whole-slide-imaging (WSI), gene expression quantification, clinical attributes like age and gender, and other biological and molecular measures [5–7]. These methods aim to quantify prognosis as a scalar hazard ratio and are supervised by Cox loss, a common method in survival analysis [8]. Effective ML-driven automated prognostication methods may reduce practitioners’ burden and resource-consumption. It should also account for challenges in data quality in low-resource areas, while helping improve patient outcomes by selecting of optimal disease management options, making it a promising active research area.

### Multimodal Machine Learning and Model Interpretability

As ML is used to tackle increasingly complex problems, recent studies have sought to leverage multiple input data modalities to make downstream predictions (i.e., multimodal modeling) rather than relying on a single data type (i.e., unimodal modeling) [9]. This follows the logic that increasing the model’s exposure to a more complete set of patient characteristics may facilitate improved predictive power by utilizing complementary information. Accordingly, many studies demonstrated the application of multimodal modeling to cancer prognostication, providing improvements over unimodal methods [10]. These works aim to study optimal approaches for fusing embeddings (i.e., vector representations) from multiple data representations - methods to combine unimodal feature representations (i.e., vectors) to capture relationships both within and between these modalities (e.g., gene expression, whole slide imaging). For example, Chen et al. [11] combined crossmodal attention gating and a Krokener-product based method to model interactions across features extracted from different input modalities (WSI, gene expression, etc.). There exist efforts to validate these technologies across cancer types and explore their interpretability to ensure they truly capture relevant biological features as means to improve reliability and avoid sources of bias.

Clinical or biological interpretability of model decision making can foster practitioner and patient trust in the technology, which is imperative for the adoption of ML technologies in the medical context [12, 13]. Several recent multimodal cancer prognostication studies have applied specific architectures and techniques to enhance model interpretability and offer explanations for model predictions [14, 15]. For instance, specialized encoding schemes which reflect prior biological structure (e.g., genes to pathways) offer opportunities to reduce noise and further increase the capacity to interrogate the model findings.

### DNA Methylation, Gene Expression, and Histopathology

Multimodal cancer prognostication methods typically utilize genomics data such as gene expression profiles or DNA methylation (DNAm). DNAm refers to the binding of a methyl group to the DNA, typically a cytosine residue of a cytosine-guanine dinucleotide (CpG), which impacts the binding of transcription factors and regulatory proteins, largely responsible for setting the transcriptional programming for specific cell-types during aging and initiating pathogenesis [16]. The proportion of methylated alleles measured at specific sites can be recorded and used as informative data. As millions of sites can be methylated, this modality is currently underutilized by cancer prognostication approaches largely due to the prohibitive dimensionality for encoding to a latent space (i.e., set of features abstracted from the input data and optimally aligned to the modeling objective). Capsule-style approaches such as sparsely coded layers (i.e., local connectivity) can circumvent these challenges and provide additional interpretability while learning meaningful representations [17, 18].

High-resolution gigapixel histopathological whole slide imaging (WSI) are informative tools in pathology workflows. As most GPU-equipped machines cannot allocate sufficient memory to apply deep learning models to the entire WSI, WSI are often broken into smaller subarrays (i.e., patches) which are modeled independently or jointly through a two-stage process. Graph neural networks (GNNs) – where nodes are representations/embeddings of patches and edge connections are formed based on spatial adjacency – are promising modeling approaches based on their ability to capture complex micro and macro architectural relationships in WSI based on spatial connectivity [19]. Few applications of GNNs to WSI for cancer prognostication currently exist [14, 20]. Further investigation of additional cancer types and experimental setups could prove beneficial.

### Pretraining in Multimodal Cancer Prognostication

Pretraining strategies are typically used as a form of regularization to tune model weights to extract meaningful representations from input data before these models are applied and trained for a specific task (e.g., similar to how face detection algorithms are finetuned on an edge device to recognize specific individuals’ faces). For example, convolutional neural networks (CNNs) are commonly pretrained for classification on ImageNet, a large dataset of natural world images with 1000 classes (ex; cats, dogs, cars). Then these models are trained on specialized objectives (ex; cancer diagnosis), having already learned to extract high level features from images (ex; shapes, edges). Recent studies have shown that multimodal neural networks – such as those integrating gene expression and DNAm – benefit from crossmodal pretraining; unimodal pretraining mechanisms which incorporate complementary modalities, such as predicting gene expression from DNAm and vice versa [21, 22]. Other promising unimodal pretraining strategies include the usage of variational autoencoders (VAEs) [23], transfer learning [24], and contrastive learning [25]. VAEs seek to reconstruct input data from low level representations, and can be beneficial when only data from one modality is available. Transfer learning centers on domain adaptation (e.g., recognizing liver tumors and adapting the model to learn to recognize Colorectal tumors) and is key in situations where a high quantity of data is available that is not directly applicable to the target task (e.g., far fewer Colon slides available as compared to liver).

Contrastive learning has emerged in recent years as an additional unimodal pretraining mechanism, where data is compared across classes or contexts (e.g., comparing spatially adjacent subarrays or the same subimage which has been augmented/re-stained) to learn important distinguishing features.

These innovative pretraining strategies have yet to be comprehensively investigated and applied to omics data such as gene expression and DNA methylation, as well as histopathology, for the purpose of prognostication. Existing works on multimodal cancer prognosis largely utilize a single unimodal pretraining method and assume its effectiveness. Unimodal networks are trained to predict survival, and their weights are subsequently transferred to encoders in the multimodal model, as it is assumed that the unimodal training was sufficient to learn relevant high level features from input data. Cheerla & Geveart [10] evaluated the effectiveness of multimodal pretraining on different cancer subtypes, investigating whether learned biology from certain subtypes could be directly applicable to other subtypes. However, they too did not investigate the usage of other specialized pretraining mechanisms in an unimodal context, such as cross modal pretraining or contrastive learning, which have shown promise in other domains but have yet to be explored for pretraining multimodal prognostication methods. Accordingly, there is a gap in the existing literature in the potential benefits of using such pretraining strategies when developing multimodal modeling methods for cancer prognostication [4].

## Contributions

Current works integrating histopathology, gene expression, and DNA methylation in a multimodal modeling approach to predict cancer prognosis can benefit from evaluating the impact of emerging pretraining strategies which may further leverage the biological and histomorphological information encoded within and across these modalities. Here, we present an application of interpretable multimodal prognostication using gene and pathway-neural networks and GNNs to compare innovative pretraining mechanisms including self-supervised crossmodal pretraining, Graph Contrastive Learning, and transfer learning. Our study is the first to jointly evaluate the usage of these methods for multimodal cancer prognostication. Crucially, we bring insight on deep learning-driven prognostication – including assessment and analysis regarding pretraining strategies, as well as biological insights derived from the interpretation of models – which can inform further work in the field. We conduct our study on publicly available data from The Cancer Genome Atlas (TCGA) from 8 cancer subtypes: [4].

## Methods

### Methods Overview

In **Table 1** and **Figure 1**, we have included a glossary and graphical overview of the methods employed in this study and their definitions for reference. In brief, the following methodologies were adopted for this work:

**Table 1:**
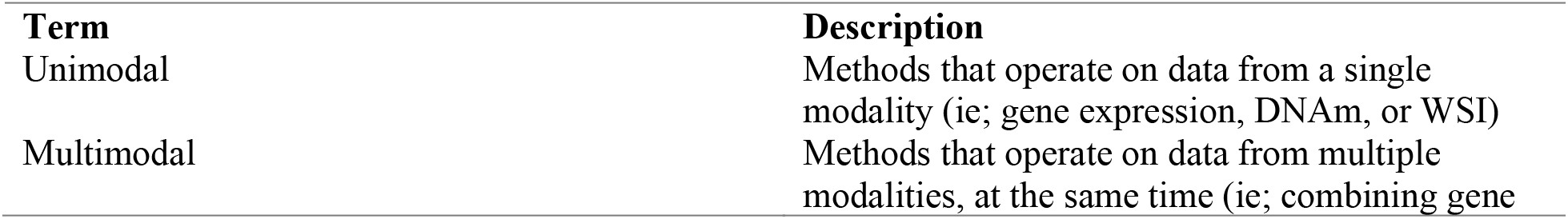
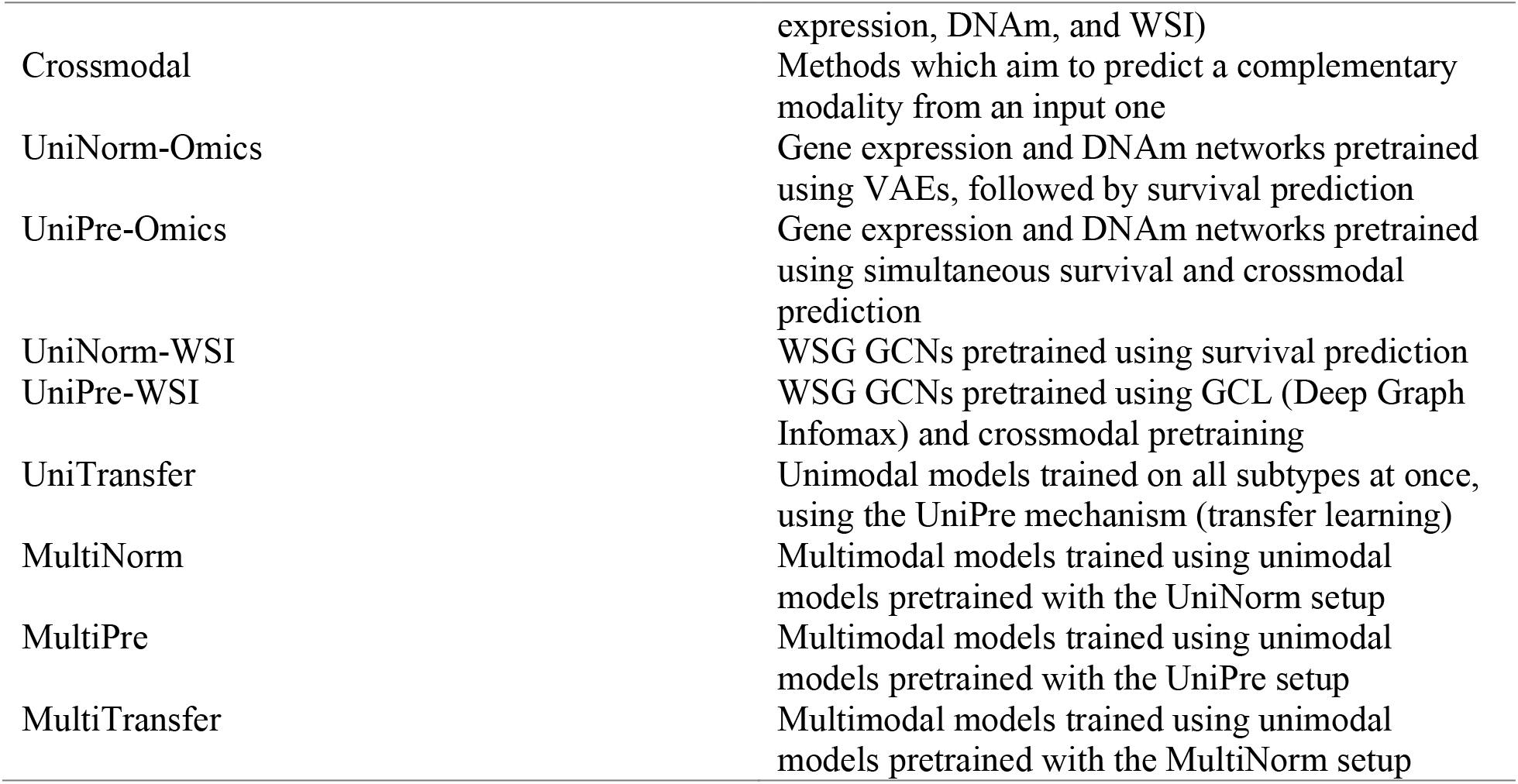
Definition of terms used in this analysis

**Figure 1:**
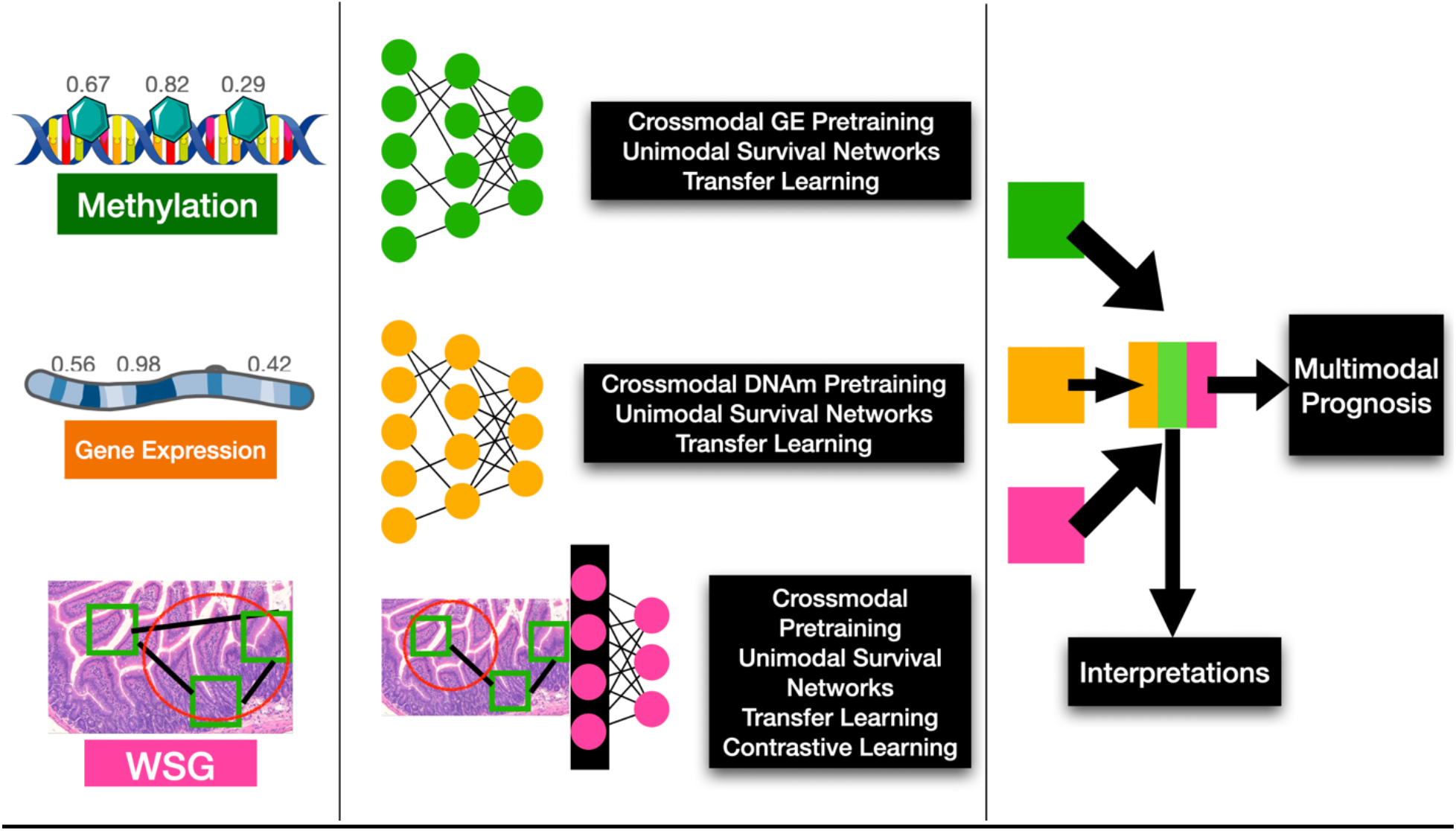
Study workflow overview; neural networks are pretrained using various methods to encode gene expression, DNA methylation, and histopathology whole slide imaging data. Multimodal prognostication models are developed using these encoders pretrained using these methods, to assess the utility of each one.

1. **Data Collection:** Multimodal data (e.g., DNAm) from the TCGA were collected and preprocessed from 8 cancer subtypes
2. **Model Approaches:** Pathway neural networks were leveraged for omics data, while graph convolutional neural networks were used to capture information across tissue slides
3. **Unimodal Pretraining:** Several pretraining strategies were compared for unimodal prediction (e.g., supervised learning, crossmodal, contrastive, transfer learning)
4. **Multimodal Pretraining:** Several pretraining strategies were compared for multimodal prediction (e.g., supervised learning, crossmodal, contrastive, transfer learning)
5. **Model Comparison:** Models were compared through reports of concordance statistics and confidence intervals. Partial likelihood ratio tests and Kaplan Meier curves were devised to compare the suitability of deep learning-derived predictors to pTNM staging, etc.
6. **Model Interpretation:** Models were interpreted using neural network interpretation methods (e.g., integrated gradients, pathway layers, graph convolutional network pooling layers), comparison to tumor-infiltrating lymphocytes and enrichment analyses to determine important genes, pathways, and WSI regions of interest

### Dataset and Preprocessing

The Genomic Data Commons (GDC) data transfer tool was used to download DNA methylation data, gene expression quantification, clinical data, and histopathological whole slide tissue images from the TCGA subtypes BLCA, BRCA, HNSC, KIRC, LIHC, LUAD, PAAD, and SKCM. We restricted to patients where gene expression quantification, DNA methylation, a WSI downsampled to 20x magnification, and clinical data were all available. Survival outcomes were right censored and recorded as days to death. Cases from each subtype were partitioned into an 80/10/10 train-test-validation split. We ensured cases within these sets exhibited similar survival status through report of a log-rank statistical test that compared survival between the dataset partitions. Gene expression and DNA methylation data were preprocessed using standard functions from the TCGABiolinks R package via Sesame [26]. Gene expression data was FPKM-normalized and log-transformed. DNA methylation and gene expression data were preprocessed for each patient by only including CpGs and genes, for which data was available for every patient in the cohort across all cancer subtypes.

Stain normalization was applied to the WSI using the PathFlowAI Python package [27] (Macenko method) to match the application of staining reagents at the host institution. PathFlowAI was used to split each image into non-overlapping 256×256 subimages (i.e., patches). These subimages were transformed into embeddings (ie; vector representations) using a ResNet50 Convolutional Neural Network (CNN) [28] encoder pretrained using the ImageNet dataset, which is a common protocol in WSI patch encoding. These subimages served as nodes of a graph which were connected based on spatially adjacency using a radius neighbors algorithm. The embeddings formed the nodal attributes. The largest connected component was selected for each whole slide graph (WSG) to represent each WSI.

Clinical characteristics (e.g., sex, pathological stage, subtypes) are presented in **Appendix Table S1**.

### Unimodal Omics Pretraining

All omics encoders consist of sparsely coded and gene/pathway-informed layers, where input nodes represent gene expression and DNA methylation. The following layers represent biological pathways and genes associated with input genes and CpGs respectively. In standard neural networks, a single node in each layer is connected to every node in the following layer. When sparse coding is applied, each node in the input layer (gene expression, DNAm) is only connected to a certain subset of nodes (aka; capsules) in the following layer (genes, pathways) (**Figure 2**). The Gene Ontology (GO) Biological Processes Set was used to construct gene expression pathway nodes (https://www.ebi.ac.uk/QuickGO/annotations), while genes associated with each CpG according to Illumina documentation are used as a gene set for DNA methylation (GSE42409). This capsule-style neural network configuration resembles the approach used by Gold et al., 2019 [29].

**Figure 2:**
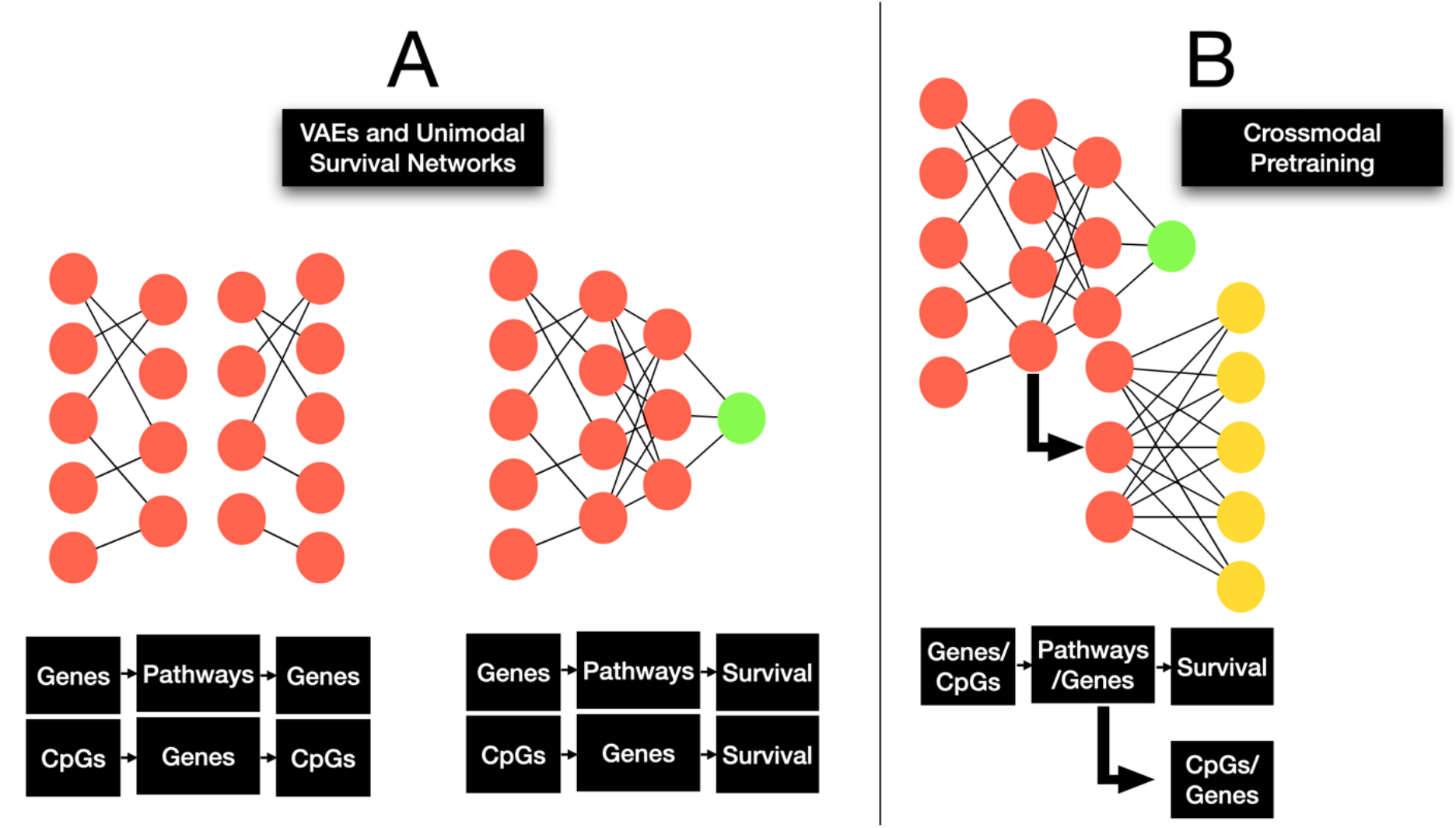
Unimodal omics pretrraining; A) UniNorm-Omics procedure consisting of unsupervised VAE pretraining followed by finetuning using a survival prediction objective; B) UniPre-Omics procedure including concurrent crossmodal pretraining and survival prediction

The following unimodal pretraining approaches were compared:

#### *Unsupervised/Supervised Pretraining* (UniNorm-Omics)

Unimodal encoders for gene expression quantification and DNA methylation data were individually pretrained for each subtype, using variational autoencoders (VAEs). VAEs are commonly used to generate compressed representations of high dimensional data, and subsequent unimodal survival training encouraged the encoders to learn features relevant to cancer prognosis while requiring less data. By transferring the initial weights for the encoders of the top-performing VAEs for each subtype, these neural networks were finetuned to predict prognosis from the omics information (UniNorm-Omics) directly.

#### *Crossmodal Pretraining* (UniPre-Omics)

Separately, neural networks were trained to simultaneously predict prognosis, and reconstruct information from complementary modality. The complementary modality for gene expression is DNA methylation beta values, and vice versa. Input data is first encoded to a common latent space, followed by one branch of hidden layers used to predict the complementary modality, and another nearly identical branch that outputs prognosis predictions. The primary advantage of our crossmodal supervision strategy (UniPre) was to learn more nuanced information relevant to other modalities, which may bring a richer understanding of input data as understanding of other modalities given a single data type can enhance the quality of learned features by forcing models to learn complex information.

VAEs were trained for a total of 500 epochs and checkpointed every 100 epochs, optimized by the Adam optimizer, and employed a learning rate of 0.008. A hyperparameter scan was performed to determine the optimal VAE checkpoint for beginning unimodal omics survival training for each cancer type, by considering VAEs checkpointed at 100, 200, 300, 400, and 500 training epochs. Sparsely coded unimodal survival networks were supervised by a Cox loss for prognostication and a Mean Squared Error objective for crossmodal prediction. These networks were trained for up to 40 epochs with a learning rate of 0.0002, a batch size of 32, a linearly decaying weight scheduler set to a decay of 1e-4, and the Adam optimizer.

### Unimodal WSI GCN Pretraining

Unimodal GCN models were trained to predict patient prognosis from WSGs. GCN models consisted of several blocks of SAGEConv graph convolutional layers and SAGEPool node pooling layers [30], which jointly learn contextualized node embeddings by pooling graphs using a self-attention mechanism where attention scores are calculated using graph convolutions.

Spatial information was aggregated across the WSG using a graph-wide JumpingKnowledge (JK) pooling layer [31] to output a graph-level embedding. This embedding is passed through several fully connected layers to obtain a scalar hazard prediction.

The following unimodal pretraining approaches were compared (**Figure 3**):

**Figure 3:**
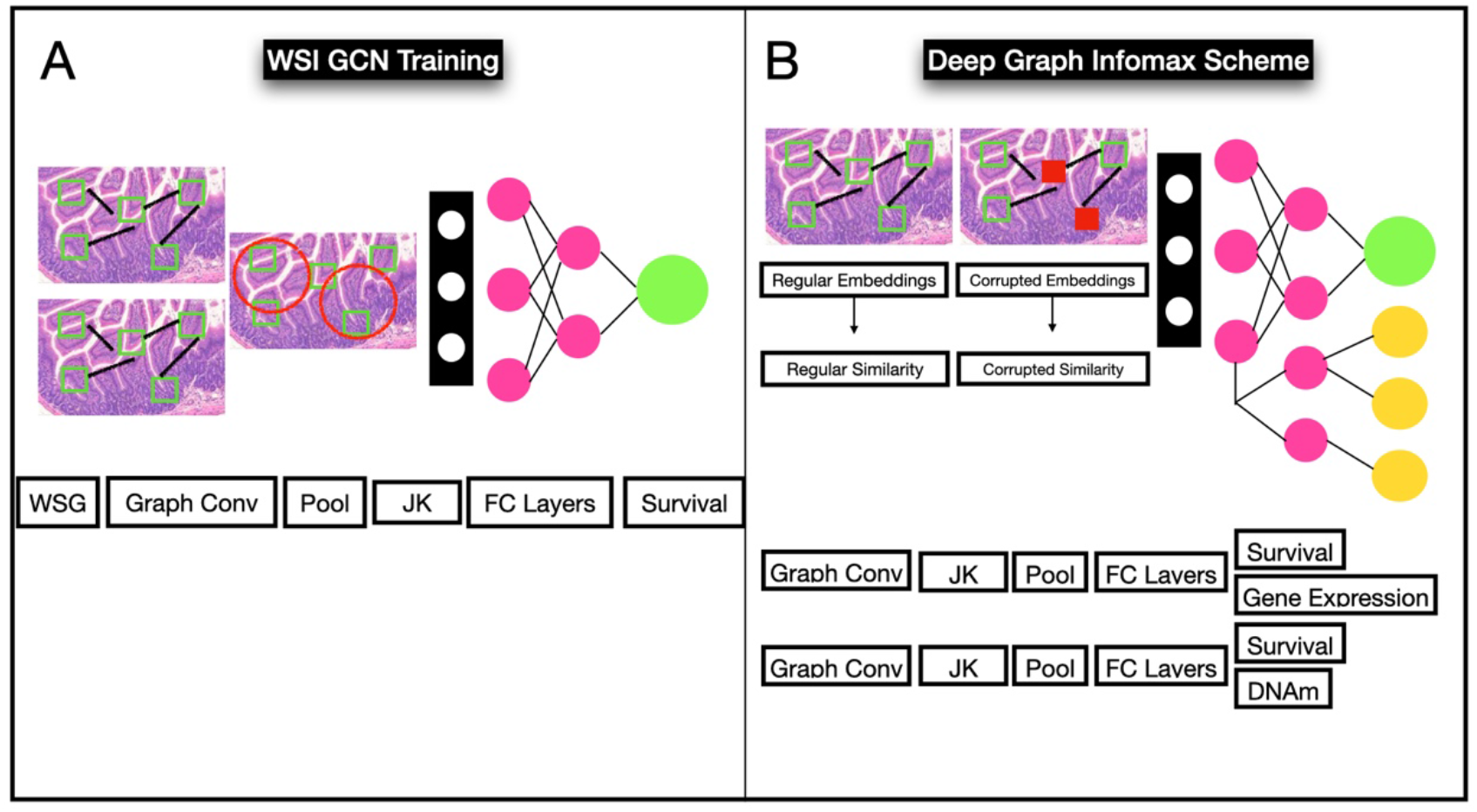
Unimodal WSI GCN pretraining; A) GCN pretrained to predict survival; B) GCN pretraining using Graph Contrastive Learning (Deep Graph Infomax scheme) followed by concurrent crossmodal pretraining and survival prediction

**Figure 4:**
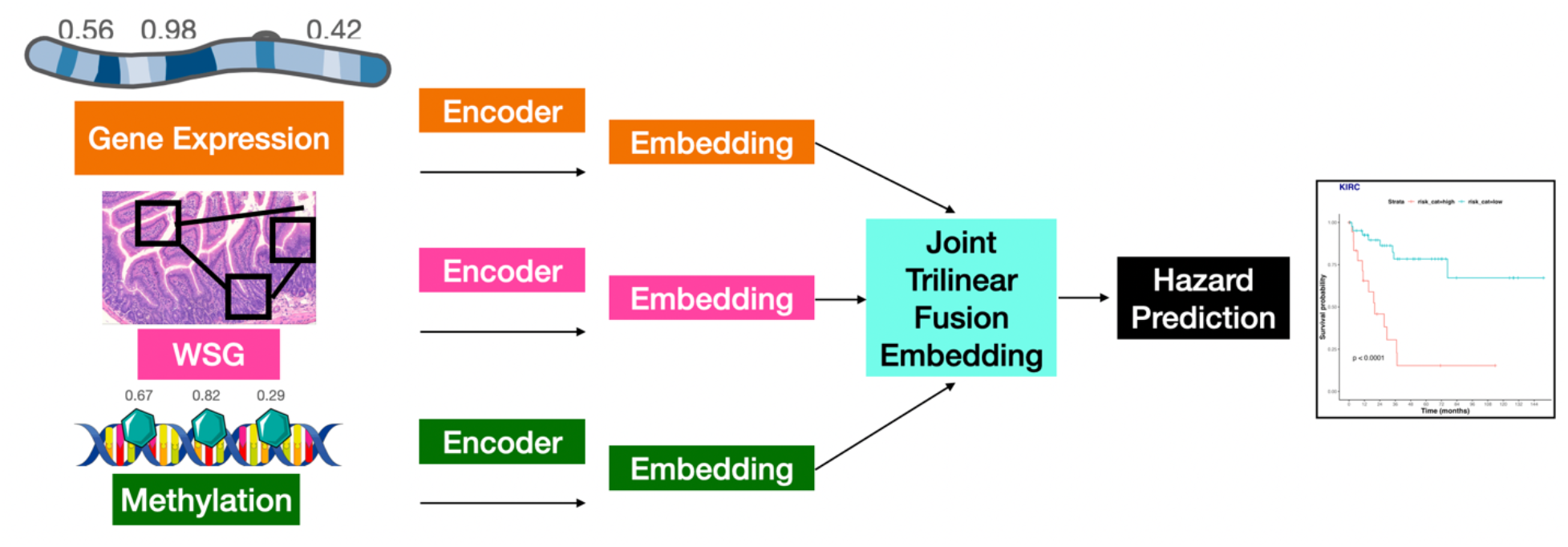
Multimodal modeling; Pretrained encoders are used to extract features from gene expression, DNAm, and WSI; embeddings are fused using Trilinear Fusion; prognosis is predicted using a feedforward neural network. Predicted hazards are dichotomized to generate a Kaplan Meier plot to compare low and high risk groups.

**Figure 5:**
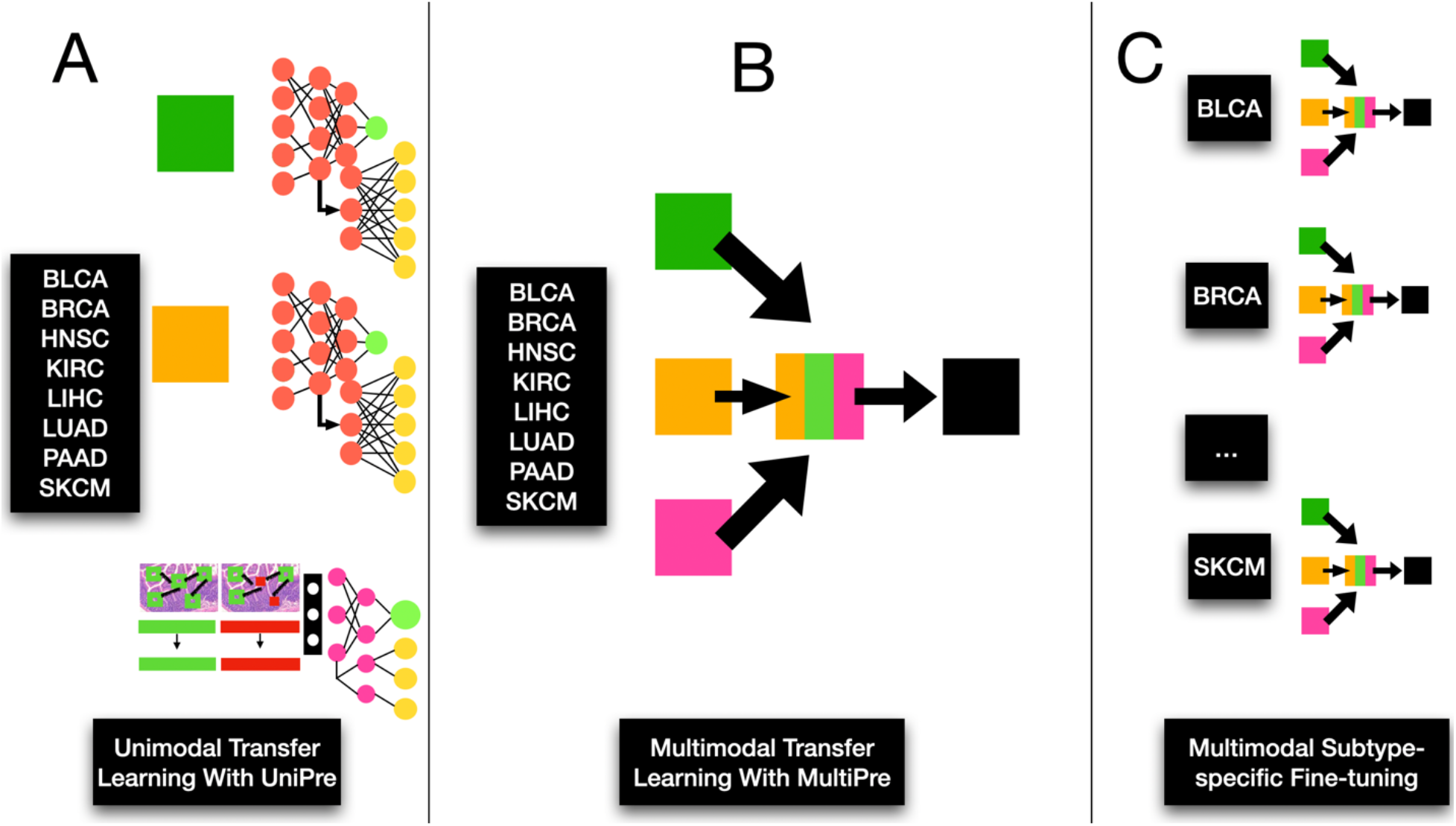
Transfer learning experimentation; unimodal transfer learning with the UniPre mechanism followed by multimodal transfer learning and subtype-specific multimodal finetuning.

**Figure 6.**
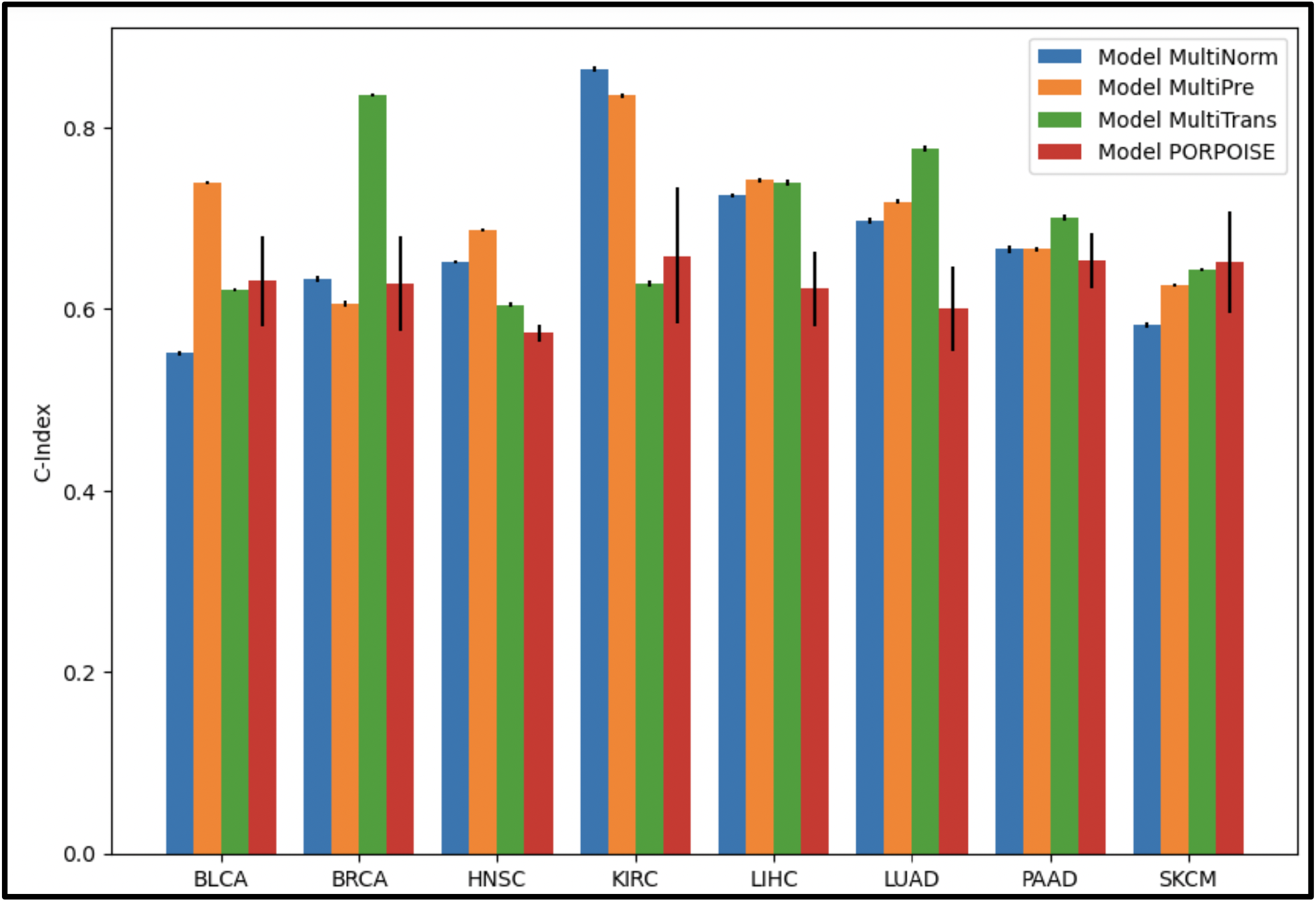
Bar graph of multimodal model performance indicated by C-Index across subtypes

#### Supervised Pretraining (niNorm-WSI)

GCNs were trained to predict prognosis using standard randomly initialized weights (**UniNorm-WSI**).

#### Crossmodal and Contrastive Pretraining (UniPre-WSI)

Separately, a crossmodal pretraining procedure was adopted as outlined in the previous section, using weights transferred from a pretraining that used graph contrastive learning (GCL) (**UniPre-WSI**). GCL pretraining was conducted by training a Deep Graph Infomax [32] scheme to update patch-level embeddings before including of the JumpingKnowledge and fully connected layers for whole-graph embeddings. Deep Graph Infomax is a commonly used GCL approach for learning node and graph embeddings. These graph level embeddings were used for simultaneous prognostication and crossmodal training through two separate neural network branches which predict prognosis and information from the complementary modality, e.g., DNA methylation or gene expression, respectively. GCNs were separately trained to predict both DNAm and gene expression in the crossmodal scheme, to assess which modality brings improved complementary information extraction for prognostication, for each subtype. The top performing model for each subtype (between DNAm and GE crossmodal pretraining) was reported as the selected crossmodal approach. GCL models were trained for 15 epochs, and weights were transferred to the subsequent prognostication GCN model from the training epoch which obtained the lowest validation loss. These models were implemented using the PyGCL Python library (Zhu et al, 2021).

GCNs for prognosis and crossmodal were trained using the Cox and Mean Squared Error losses respectively, with a coarse hyperparameter search over the following learning rates for each cancer subtype: 1e-4, 2e-4, or 4e-4. All networks used the Adam optimizer with a linearly decaying weight scheduler set to 1e-4. Models were trained with a batch size of 4 due to GPU memory constraints, and gradient accumulation every 4 steps to account for the small batch size (effective batch size of 16).

### Multimodal Modeling

Multimodal prognostication models were developed for each cancer subtype. These included the top-performing unimodal models for each modality (as identified using the validation Cox loss), from which features were extracted from the penultimate layer. Features from all modalities were combined using a gated attention and Krockener product-based tensor fusion mechanism introduced by Chen et al [11], as this method allows for the learning of heterogeneous features across modalities. The fused representation is passed through several fully connected layers, followed by a final prognosis prediction output layer. Two sets of multimodal models were trained for each subtype:

1. **MultiNorm**– similar to **UniNorm-Omics** and **UniNorm-WSI**, which initializing weights from networks trained using these schemes (e.g; VAEs and unimodal survival training), and
2. **MultiPre–** which additionally employed the pretraining strategies (e.g., unsupervised, crossmodal, contrastive) and initialized weights from **UniPre-Omics** and **UniPre-WSI**.

This allowed us to compare the effectiveness of the optimal pretraining strategies when predicting prognosis in a multimodal context.

Multimodal models were trained with a learning rate of 0.0001 for up to 40 epochs, using the Adam optimizer with a linearly decaying weight scheduler set to a value or 1e-4, and a batch size of 3 due to GPU memory constraints. Gradient accumulation was performed every 8 steps to account for the small batch size (effective batch size of 24). Unimodal encoders were frozen for the first 10 training epochs to combat overfitting. Training was supervised by the Cox loss.

### Cross-Subtype Transfer Learning Experimentation

Using cross-subtype transfer learning, Cheerla & Geveart [10] found that it may be beneficial to apply learned biological information from one or more subtype(s) to prognostication of a different subtype, potentially due to the presence of shared biology from similar tissue contexts [33]. We used this finding to motivate our assessment of this multimodal pretraining approach. Transfer learning was conducted by first training a single model per modality, using the UniPre-procedure of concurrent crossmodal pretraining and survival prediction (**UniTransfer-Omics**), as well as GCL for WSI (**UniTransfer-WSI**). For WSI GCN, the gene expression prediction scheme was used for crossmodal pretraining, as it performed better than the DNAm scheme for non-transfer learning UniPre models. In this instance, these models were trained on patients across all subtypes to limit the number of comparisons (i.e., a single subtype per model). A multimodal (**MultiTransfer**) prognostication model which used starting weights from the transfer learned-unimodal models, was similarly trained on data from all subtypes for 10 epochs. Finally, the **MultiTransfer** model was separately finetuned on each subtype for 40 epochs with a learning rate of 1e-3 (after coarse hyperparameter search), and evaluated.

### Baseline Clinical Experimentation

Random-forest-based models and a Cox Proportional Hazards (CoxPH) model, were trained per subtype to predict prognosis from clinical data. These were implemented with the Sksurv Python package [34], and trained using standard clinical covariates including age, race, and sex.

Additionally, a separate set of random-forest based prognosis models were trained which included pathological stage (pTNM) as an additional covariate. The clinical models served as a baseline to compare unimodal and multimodal methods against.

### Evaluation and Interpretation Experiments

#### Model Performance

Prognostication performance was measured using the concordance index (C-index), calculated on survival predictions on the held-out testing set for each subtype. We reported the C-index for unimodal prognostication models (**UniNorm-, UniPre-**, and **UniTransfer**), clinical forest-based methods, and multimodal models (**MultiNorm, MultiPre**, and **MultiTransfer**). A 1000 sample non-parametric bootstrapped 95% confidence interval C-index was calculated for each model. Multimodal models are also compared against reported results for PORPOISE [15], the current state of the art for interpretable multimodal cancer prognostication, as well as Cheerla & Gevaert [10], a prominent pioneering work in multimodal prognostication though their study does not center on interpretability. An evaluation was not performed on identical testing folds, so one-to-one comparisons were avoided.

#### Kaplan Meier and Statistical Comparisons

Hazards were dichotomized across the top-performing multimodal models to generate Kaplan-Meier (KM) survival curves along with Log-Rank testing conducted on the non-training folds of each subtype. High-risk and low-risk groups were delineated using sensitivity analysis on predicted hazards for each subtype (i.e., a threshold determined via a sensitivity analysis of a log-rank statistic and standardized regression coefficients from CoxPH models fit on dichotomized hazards). This was performed using the R package ‘survminer’ [35] and the ‘surv_cutpoint’ function. We also used partial likelihood ratio testing to compare CoxPH models fit on the predicted hazards and clinical data versus the clinical data alone to assess whether incorporating deep learning hazards from multimodal models alongside clinical data could improve survival curves compared to the usage of solely clinical data, and to assess the added benefit of hazards predicted using our multimodal models versus the sole usage of advanced pTNM to predict high/low risk (p-value less than 0.05 indicates improved prognostic capacity; less than 0.1 is suggestive) [36].

#### Gene/Pathway Interpretation

We additionally interrogated these models using layer-wise Integrated Gradients from the Captum library [37, 38] to report the top 10 most significant genes (from DNA methylation capsules) and biological pathways (from gene expression capsules) per subtype. Integrated Gradients is a method that can identify significant genes/pathways by integrating the gradients (with respect to the genes/pathways) accumulated from a reference (i.e., low risk) to target group (i.e., high risk). In this experiment, the low risk patients were used as a reference, while high risk patients were the target group. Genes and pathways were ranked using this method. Pathway analysis was conducted using Enrichr, to interrogate significant genes from the DNAm capsules to evaluate their potential biomarker applicability. Fused unimodal embeddings from were extracted from the MultiTransfer model prior to finetuning on subtypes by passing input data through the network through the trilinear fusion layer, and visualized using the t-distributed Stochastic Neighbor Embedding (t-SNE) method. Additionally, we sought to provide evidence that these models are able to provide predictive value above and beyond established molecular subtypes which have been implicated in prognosis. As an example, we restricted the KIRC cohort to individuals with BAP1 and PBRM1 mutations (prior studies have identified their association with survival). Oncoprint queries for these KIRC molecular subtypes and copy number alteration (CNA) data were downloaded to facilitate this analysis. Partial likelihood ratio testing and generalized linear modeling was used to assess whether predicted hazards provides additional predictive value versus the BAP1 and PBRM1 subtypes of KIRC[39]. KM plots were generated conditional on each KIRC molecular subtype to demonstrate the added predictive value.

#### Alignment of Important Regions of Interest in WSI with Immune Infiltration

Finally, we visually inspected areas of tissue from randomly selected slides which were assigned high importance by the WSI GCN as a basis of explanation for features of histopathology relevant for multimodal prognostication. Patches present in the WSG following the final SAGEPooling layer were determined to have high importance. These regions were compared against corresponding published maps of tumor infiltrating lymphocytes (TILs) [40]where such information was made publicly available. TILs may confer prognostic information in cancer imaging, as the density, type, and spatial localization of TILs have been shown to be associated with prognosis [41].

Thus, we hypothesized that capable models would assign relatively high importance to tissue areas containing TILs, especially for predicting low risk patients. Wald-Wolfowitz testing [42] was used to assess agreement between TIL maps and high-attention tissue regions, where the null hypothesis was that TILs and high-attention image patches were in high agreement, i.e., drawn from the same distribution. For each subtype included in this analysis, a p-value was generated using Wald-Wofowitz testing for each slide for which a corresponding TIL map was available [43]. A higher p-value indicates greater TIL localization by the GCN model, while a lower p-value indicates the opposite. Fisher testing was used to determine the significance, direction and magnitude of the relationship between model’s localization of TILs and predicted hazards (as determined through the aforementioned dichotomized hazards; i.e., how responsible are TILs for model’s hazard predictions). Separately, Kaplan Meier plots were generated for all available patients using TIL localization as further stratification between high and low risk groups to demonstrate the capacity for improved predictiveness beyond the predicted hazards. Partial likelihood ratio testing was used to corroborate with the KM analysis (i.e., does the alignment of the model to TILs provide additional predictive value). KM plots stratifying patients by high and low concordance between the model and TIL maps were generated, separately for patients of high and low predicted risk. Likewise, KM plots stratified by risk were generated, separately for patients by high and low concordance between the model and TIL maps. We retained KM plots suggestive of an interaction between the TIL phenotype and risk strata. This analysis is reported for subtypes for which TIL maps were available: BLCA, BRCA, LUAD, PAAD, and SKCM.

Reported significant genes, pathways, and image region visualizations were inspected by a certified pathologist to validate their potential applicability as biomarkers.

## Results

### Unimodal Prognostication Performance

Unimodal models were trained to predict survival using standard pretraining procedure (UniNorm), crossmodal pretraining (UniPre), and transfer learning (UniTransfer). Separately, clinical models were trained to predict survival using age, race, sex, and both with/without pathological stage.

In **Table 2** we report the 95% confidence interval bootstrapped C-indices for each subtype for the UniNorm models, and we report the same statistic for the UniPre models in **Table 3**. These mechanisms are compared in **Appendix Figure S2** as well. Results for the UniPre-WSI model, when DNAm was used as the crossmodal target, are reported in **Appendix Table S3**.

**Table 2:** Unimodal boostrapped C-indices with 95% confidence intervals for UniNorm models

| Subtype | DNAm | Gene Expression | WSI GCN |
| --- | --- | --- | --- |
| BLCA | 0.633 ± 0.002 | **0.689 ± 0.002** | 0.564 ± 0.003 |
| BRCA | **0.711 ± 0.002** | 0.547 ± 0.002 | 0.558 ± 0.003 |
| HNSC | 0.600 ± 0.002 | 0.608 ± 0.002 | **0.680 ± 0.002** |
| KIRC | 0.699 ± 0.003 | 0.691 ± 0.005 | **0.777 ± 0.004** |
| LIHC | 0.562 ± 0.003 | 0.569 ± 0.002 | **0.722 ± 0.003** |
| LUAD | 0.645 ± 0.003 | 0.601 ± 0.002 | **0.663 ± 0.003** |
| PAAD | 0.603 ± 0.004 | 0.585 ± 0.004 | **0.616 ± 0.004** |
| SKCM | 0.588 ± 0.003 | **0.632 ± 0.003** | 0.623 ± 0.002 |

**Table 3:** Unimodal bootstrapped C-indices with 95% confidence intervals for UniPre models

| Subtype | DNAm | Gene Expression | WSI GCN |
| --- | --- | --- | --- |
| BLCA | 0.619 ± 0.003 | **0.692 ± 0.002** | 0.571 ± 0.002 |
| BRCA | 0.552 ± 0.003 | **0.624 ± 0.002** | 0.573 ± 0.003 |
| HNSC | 0.623 ± 0.002 | **0.648 ± 0.002** | 0.606 ± 0.003 |
| KIRC | 0.688 ± 0.002 | 0.713 ± 0.002 | **0.794 ± 0.004** |
| LIHC | 0.618 ± 0.003 | 0.655 ± 0.003 | **0.746 ± 0.002** |
| LUAD | 0.646 ± 0.002 | 0.631 ± 0.004 | **0.669 ± 0.003** |
| PAAD | 0.604 ± 0.004 | **0.628 ± 0.004** | 0.575 ± 0.004 |
| SKCM | 0.590 ± 0.003 | **0.640 ± 0.003** | 0.559 ± 0.002 |

Performance of baseline clinical models (trained using age, race, sex, with/without stage) is presented in in **Appendix Table S4**. C-index performance for UniTransfer models is reported in **Appendix Table S5** (not finetuned on each subtype).

In the absence of pretraining, WSI GCNs outperform DNAm and gene expression unimodal models on majority of subtypes with the UniNorm models, while gene expression becomes the top-performing modality on more than half of subtypes when crossmodal pretraining is implemented with the UniPre models. All UniPre gene expression models outperformed the corresponding UniNorm models (6.62% average increase). UniPre DNAm models brought performance improvements on majority of subtypes compared to the corresponding UniNorm models (increase average 2.89%). UniPre WSI GCNs brought improvement on majority of subtypes compared to the UniNorm setup (increase average 2.07%). Every subtype benefitted from crossmodal pretraining for at least one modality. Top performing UniNorm or UniPre models outperformed top-performing clinical models which did not include stage for all subtypes (increase average 34.05%), and outperformed clinical models which did include stage (increase average 12.44%), for all subtypes besides BLCA.

### Multimodal Prognostication Performance

In **Table 5**, we report the test set 95% confidence interval bootstrapped C-Indices for the MultiNorm, MultiPre, and MultiTransfer models. We also add comparisons for analogous models developed by the PORPOISE study and Cheerla & Gevaert. Furthermore, in **Figure 7**, we present Kaplan Meier survival curves for each subtype, generated using the top performing multimodal model for the given subtype. In **Appendix Table S6**, we report the partial likelihood ratio testing results of the partial likelihood ratio testing to assess whether the inclusion of both clinical covariates and deep learning hazards improved the predictiveness of clinical covariates or deep learning hazards alone. These results indicated that this was the case for most cancer subtypes, where deep learning provides improved prognostic capacity beyond clinical staging and vice versa (i.e., clinical staging used in conjunction with the deep learning model provides added predictive value).

**Table 5:** Comparison of multimodal models via their C-indices and 95% confidence interval; * No reported confidence interval

| Subtype | MultiNorm | MultiPre | MultiTransfer | PORPOISE | Cheerla and Gevaert * |
| --- | --- | --- | --- | --- | --- |
| BLCA | 0.551 ± 0.003 | **0.739 ± 0.002** | 0.621 ± 0.002 | 0.631 ± 0.050 | 0.73 |
| BRCA | 0.633 ± 0.003 | 0.606 ± 0.004 | **0.836 ± 0.002** | 0.628 ± 0.053 | 0.79 |
| HNSC | 0.651 ± 0.002 | **0.687 ± 0.002** | 0.605 ± 0.002 | 0.573 ± 0.010 | 0.67 |
| KIRC | **0.864 ± 0.003** | 0.835 ± 0.003 | 0.628 ± 0.003 | 0.659 ± 0.075 | 0.73 |
| LIHC | 0.725 ± 0.003 | 0.742 ± 0.003 | 0.739 ± 0.003 | 0.622 ± 0.042 | **0.77** |
| LUAD | 0.697 ± 0.003 | 0.718 ± 0.002 | **0.776 ± 0.003** | 0.600 ± 0.046 | 0.73 |
| PAAD | 0.666 ± 0.004 | 0.666 ± 0.003 | 0.700 ± 0.003 | 0.653 ± 0.030 | **0.74** |
| SKCM | 0.582 ± 0.003 | 0.626 ± 0.002 | 0.643 ± 0.002 | 0.651 ± 0.056 | **0.72** |

**Figure 7:**
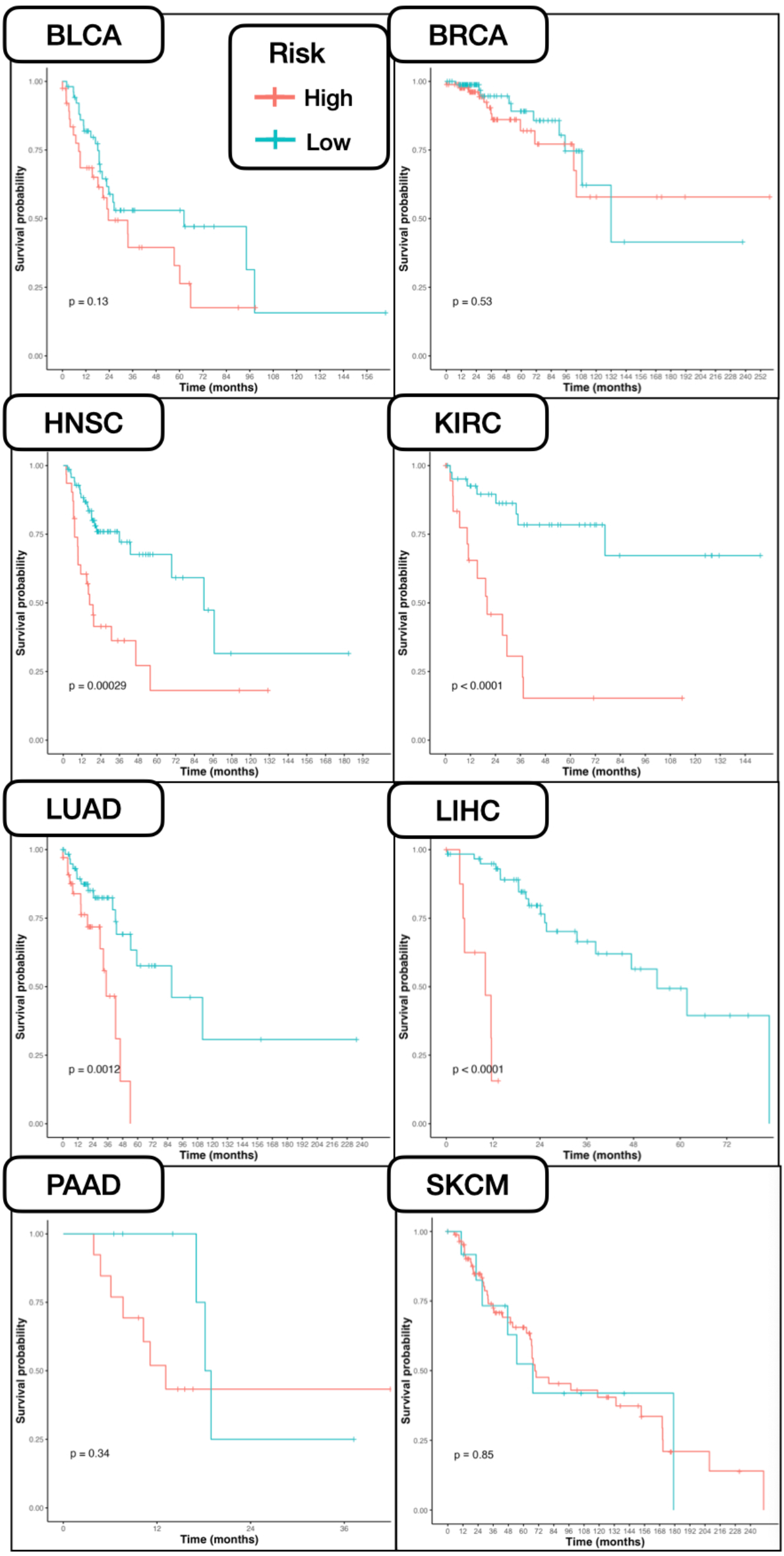
Kaplan Meier survival curves generated using top performing multimodal models on held out data.

The prognostic performance of multimodal models was improved when applying pretraining mechanisms with the MultiPre or MultiTransfer protocols rather than the standard MultiNorm setup, for every subtype besides KIRC. Top-performing models for each subtype further appear to outperform similar multimodal models presented in the state-of-the-art PORPOISE study (average C-index increase 11.54%), for every cancer type other than SKCM. These models outperformed prior pancancer-pretrained models from Cheerla & Gevaert for 5/8 subtypes (average C-index increase 6.85%).

### Interpretation

In **Appendix Tables S7** and **S8**, for each subtype, we present significant pathways and genes derived from the gene expression and DNAm encoders of top-performing multimodal models. **Appendix Table S9** reports the results of pathway analysis conducted on significant genes from the DNAm multimodal encoder. From the gene expression capsules, we identified several pathways associated with disease progression– for instance: 1) regulation of smooth muscle cell proliferation (BLCA) [44], 2) mammary gland development (BRCA) [45], 3) establishment of skin barrier (HNSC) [46], 4) regulation of NF-KB signaling (KIRC) [47, 48], 5) negative regulation of production of molecular mediator of immune response (LIHC) [49], 6) positive regulation of granulocyte chemotaxis (LUAD) [50], 7) positive regulation of cell population proliferation (PAAD) [51], and 8) epidermis development (SKCM) [52], amongst others. From the DNAm capsules, we identified the following pathways: 1) glucocorticoid receptor pathway (BLCA) [53], 2) G2 phase pathway (BRCA) [54], 3) Y branching of actin filaments (HNSC) [55], 4) regulation of heat shock proteins (KIRC) [56], 5) autophage and lysosome dysregulation (LIHC) [57], 6) semaphore interactions (LUAD) [58], 7) E2F transcription factors (PAAD) [59], and 8) PIP biosynthesis (SKCM) [60]. **Appendix Figure S10** presents t-SNE embeddings from MultiTransfer prior to subtype finetuning to demonstrate both the delineation of subtypes during pretraining and the separation of survival. Cancers with similar prognosis are generally clustered together, while those with differing prognostic profiles have more distinct clustering patterns with respect to one another. For example, LIHC has a relatively poor prognosis for most patients [61], while BRCA and KIRC have higher survival rates [62, 63]. Accordingly, BRCA and KIRC are generally clustered together, while LIHC is more distinct from the two.

Partial likelihood testing used to assess whether the hazard prediction provides additional predictive value over prognostically associated BAP1 and PBRM1 mutations (performed to investigate KIRC underperformance with MultiPre and MultiTrans), which yielded p-value of 0.041, indicating predictive improvement versus relying on the KIRC-subtyping alone (KIRC-subtype+hazard>KIRC-subtype). Combining the hazard prediction with the mutational status did not provide added predictive value over using the hazard alone (p=0.245; KIRC-subtype+hazard>hazard). From the Cox-PH models, hazard was a predictor of time to death for BAP1 patients with a p-value of 0.0739, while it predicted time to death for PBRM1 patients with a p-value of 0.000458. When controlling for subtype, hazard is predictive of death with a p-value of 1.29e-5. The KM plots in **Figure S11** suggests that the hazard provides predictive value of death, independent of key KIRC molecular alterations.

We demonstrate regions of interest (ROI) from select tissue slides which were assigned high importance after pooling by the WSI GCN from top-performing multimodal models, and compare these high-attention patches with TIL maps (**Figure 8**). Inspection of randomly selected slides and fisher’s exact testing indicated that the assignment of hazards across multiple cancer subtypes was associated with the alignment of important ROI with positions of TILs in these slides. In **Appendix Table S12**, we report the significance of TIL localization indicating lower patient risk. In **Appendix Figure S13**, we present Kaplan Meier survival curves generated using dichotomized hazards from top-performing multimodal models, which were further substratified by cases with TIL agreement with the multimodal model, for all patients for whom TIL maps were available. For SKCM, the distance between survival curves for patients with a TIL phenotype as determined by the machine learning model, was much greater than for patients without the TIL phenotype (**Appendix Figure S13A**). Likewise, the TIL phenotype as identified by the deep learning model was predictive of favorable prognosis for patients already stratified by low hazard (**Appendix Figure S13B**), suggesting an interaction between model predicted hazards and model identified TIL phenotype.

**Figure 8.**
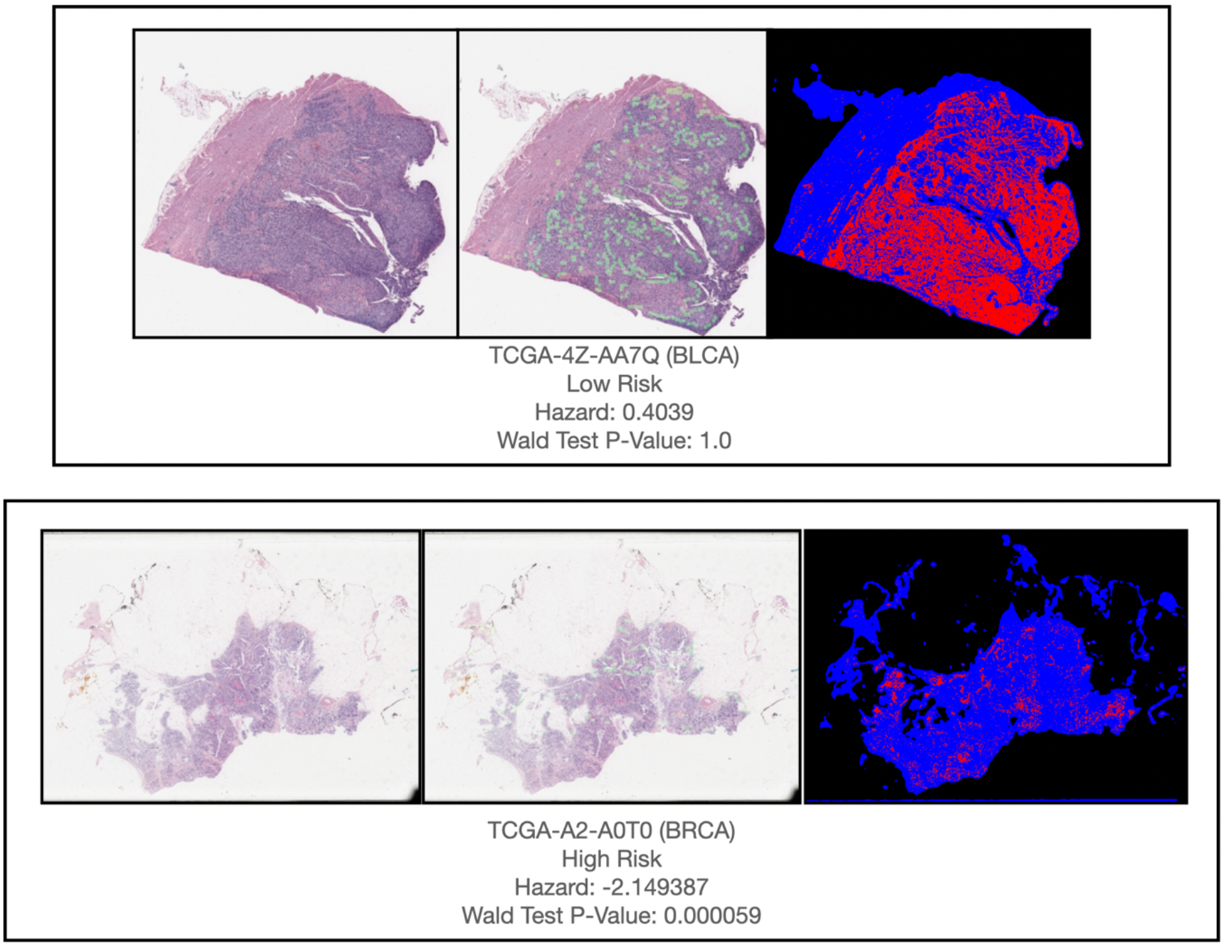
Regions of high importance (patches present after final pooling layer) from WSI GCNs of top-performing multimodal models, visualized across randomly selected held-out slides

## Discussion

Evaluation of patient prognosis provides key insight for health practitioners when considering multiple disease management options. Interpretable multimodal deep learning approaches have shown time and time again the remarkable capacity to infer prognosis from heterogenous data modalities by elucidating salient complementary features. Here, we aimed to compare and investigate emerging pretraining strategies to improve the application of these methods.

Prior works have not evaluated these nascent strategies– crossmodal pretraining, contrastive learning, and transfer learning, for unimodal and multimodal pretraining; despite initial investigations of multimodal transfer learning. Here, we demonstrate that implementing these methods on DNA methylation, gene expression, and whole slide imaging data leads to improvements in prognosis prediction compared to baseline clinical methods as well as the current state of the art. Our results show that leveraging effective pretraining approaches may be as critical to consider as improving the design of these neural network architectures. Thus, exploration of pretraining strategies should complement work to develop improved prognostic deep learning methods through novel architectures.

### Effectiveness of pretraining strategies

We demonstrated that superior unimodal prognostication performance can be achieved when implementing pretraining strategies for all eight subtypes included in the study. The strategies employed in this paper improved cancer prognostication for more than half of the cancer subtypes as compared to other state-of-the-art multimodal approaches. The pretraining strategies employed in this work improved multimodal prognostication performance for all but one cancer subtype (KIRC). This may be because molecular subtypes of KIRC which are predictive of prognosis, are solely identifiable with copy number alteration and mutational information which are not well recapitulated with the selected modalities. Partial likelihood testing revealed that when considering the PBRM1 and BAP1, which are two KIRC subtypes (Brugarolas, 2014) which are heavily implicated in prognosis, deep learning-predicted hazard was predictive of death independent of the mutational status. This was despite a relatively small sample size. Other molecular subtypes (ex; CIMP, PAM50, MMR) may be responsible for prognostic patterns as well. Superior performance of KIRC models as compared to the prior start of the art can be attributed to both the unique combinations of selected modalities (gene expression, DNAm, WSI) and modeling architectures selected. As transfer learning leveraged information from all of the cancer subtypes, this would suggest that multimodal features and interactions relevant to the prognosis of KIRC cancers may be distinct, at least within the subtypes leveraged for this study. BRCA showed the highest performance increase when transfer learning was used as compared to the other employed training strategies, suggesting the ubiquity of the identified pathways with other cancers included in the study. This may be due to the overrepresentation of BRCA cases in the study cohort. Although one-to-one comparisons cannot be made, our prognostication performance results appear to be competitive or superior than those presented in the state-of-the art PORPOISE study, for seven of the eight subtypes; our top performing models for each subtype outperform the PORPOISE multimodal fusion models by C-index with an 11.54% average increase per subtype. These models also outperformed Cheerla & Gevaert’s pancancer pretrained models on more than half of the subtypes. Our results also indicate that learning histopathological features associated with DNAm and gene expression (and vice versa for the other data types) during unimodal pretraining can help identify prognostic features. This result is corroborated by findings from studies applying similar techniques to different domains [64, 65]. Furthermore, we present the first application of Graph Contrastive Learning for prognosis prediction from WSI with our UniPre mechanism, and demonstrate its potential utility, as UniPre GCNs had higher C-indices than UniNorm GCNs for majority of the cancer types. Moderate-high separation was observed in Kaplan Meier curves for multimodal models from all subtypes besides SKCM, indicating overall strong multimodal performance in all but one of the subtypes. Reported statistical testing indicates that incorporating deep learning hazards from our developed multimodal models with clinical covariates enhance risk prediction using CoxPH models, compared to solely using hazards, pTNM stage or clinical covariates alone.

### Model interpretations corroborate with known metastasis targets

Using of capsule-inspired neural network designs in our framework improved the explainability of our approach. Reported significant genes and pathways, and interrogated histopathology regions given high importance, may serve as novel biomarker candidates, demonstrating the potential utility of deep learning for clinical application. As an example, lysosomes, the final component of autophagy [66], was implicated with disease progression from pathways mined from the DNAm and gene expression capsules for Hepatocellular carcinoma. As lysosomes play an important role in antigen presentation, cellular adhesion and migration, energy metabolism, setting the stage for further invasion, etc. their involvement in metastasis is supported by the findings in this work. Across multiple subtypes, our neural network models identified key hallmarks of cancer– e.g., disruption of cellular senescence and downregulation of genes that inhibit mitosis after cell damage was a common theme. As expected, T-cell receptor signaling was heavily implicated across many subtypes for both molecular assays and were important histological findings. For instance, comparison of significant tissue regions identified by our model with published TIL maps affirmed the all-important role of the tumor immune microenvironment (TIME) for informing the coordinated immune response to nodal and distant metastasis. Statistical testing confirmed that this relationship is especially present for BLCA, BRCA, and PAAD, while TIL localization may be important for high risk LUAD patients. These results makes sense in a biological context as prior studies have shown that TIL arrangement and presence are associated with prognosis [41, 67]. Yet, the inability of the WSI graph neural networks to pinpoint lymphoid and myeloid infiltrates in other tumor types can partially be attributed to the synthesis of information from DNAm, which can better represent immune component of the tumor microenvironment more so than through examination of the tissue slide. Future works can further interrogate these significant regions (e.g., tumor immune microenvironment– TIME) for specific pathological endpoints, such as whether the tumor had metastasized without direct evaluation of nodal involvement. As highly accurate cell type deconvolution approaches are able to decipher the TIME into its constituent components, integrating cellular populations inferred through DNAm presents an interesting area of follow-up. Follow-up clinical studies can also explore whether findings from deep learning models can be truly indicative of patient outcomes, and whether targeting these biomarkers (eg; significant genes from DNAm networks) lead to improved cancer therapies or whether specific strata may be more responsive to specific therapies. Results should be interpreted in context as it is important to note that these genes and pathways are considered significant after adjusting for complementary modalities (e.g., for DNAm, adjusting for / stratifying by gene expression data and histomorphological features).

### Limitations and Future Directions

A key limitation of this study is the usage of only 8 cancer subtypes. This may have limited visible improvements with transfer learning performance, as some cancer subtypes included in the study may share important prognostic features with other subtypes not present. Furthermore, heterogeneity is tied to distinct tissue-specific cellular differences and molecular subtypes within the selected solid tumor types that should be accounted for. There may also be untapped benefits from further preprocessing input data or conducting data preprocessing, such as including only highly variable genes and CpGs. Methods of color normalization for imaging as scanning quality and protocol of TCGA imaging is also highly variable. Additionally, hyperparameters were selected by coarse optimization due to model training time and resource constraints, while refined tuning procedure and architecture exploration may improve model training. The model architecture was not thoroughly explored, as it was outside the scope of the study. Clinical application of models like ours may also be hampered by the inability to account for potential missing data situations, as data from all modalities will not always be available in every scenario. We included patients for whom data is available from all central modalities, while this if often not the case in real world settings.

Furthermore, TCGA is not highly reflective of a real-world prospective cohort study, as case-control cohorts have collected samples. Future works will aim to address these limitations, by exploring new methods of WSI image patch encoding (ex; crossmodal transformers, multimodal coattention), explanation techniques, different sets of biological pathways and capsules, survival prediction from different data (ex; tumor microenvironment), stress testing such as leaving out certain data modalities, and additional pretraining methods such as student-teacher networks, distillation, and learning from spatially resolved data, which will improve the resolution of these findings and their interpretation.

## Conclusion

In this study, we showcase the utility of applying sparsely coded layers and GCNs for omics and histopathology analysis in multimodal machine learning while critically demonstrating that applying crossmodal pretraining and contrastive learning can benefit prognosis prediction from similar models and modalities. The implemented models trained with pretraining strategies appear to outperform the existing state-of-the art method on the majority of investigated cancer subtypes, and baseline clinical models. Future works will expand the scope of the work by investigating further model innovations and training strategies, and evaluation of in-house datasets.

## Availability of data and materials

The dataset supporting the conclusions of this article are available in the The Cancer Genome Atlas (<u>TCGA</u>) repository, doi:10.1038/ng.2764, doi:10.1016/j.cell.2018.03.035.

## Authors’ contributions

ZA- conceptualization of study, data analysis, figure generation, initial draft writing. AS- data analysis, figure generation. JL- conceptualization of study, project oversight. All authors participated in the writing and editing of this manuscript.

## Acknowledgements

This work was supported by the Dartmouth Hitchcock Medical Center Department of Pathology and Laboratory Sciences through the Emerging Diagnostic and Investigative Technologies program.

## List of Abbreviations

ANN: Artificial Neural Networks
C-index: Concordance index
CNA: Copy Number Alteration
CNN: Convolutional Neural Network
CoxPH: Cox Proportional Hazards
CpG: Cytosine-Guanine Dinucleotide
DNAm: DNA Methylation
GCL: Graph Contrastive Learning
GCN: Graph Convolutional Network
GE: Gene Expression
GO: Gene Ontology
GPU: Graphics Processing Units
JK: Jumping Knowledge
KM: Kaplan Meier
ML: Machine Learning
TCGA: The Cancer Genome Atlas
TIME: Tumor Immune Microenvironment
TIL: Tumor Infiltrating Lymphocytes
TME: Tumor Microenvironment
t-SNE: T-Distributed Stochastic Neighbor Embedding
WSI: Whole Slide Images

