## Supplementary Materials for "Assessment of Emerging Pretraining Strategies in Interpretable Multimodal Deep Learning for Cancer Prognostication"

### Supplementary Information

S1: Dataset clinical characteristics

| | BLCA | BRCA | HNSC | KIRC | LIHC | LUAD | PAAD | SKCM | Overall |
|---|---|---|---|---|---|---|---|---|---|
| n | 412 | 1098 | 528 | 537 | 377 | 585 | 185 | 470 | 4192 |
| age (mean (SD)) | 68.13 (10.61) | 58.63 (13.20) | 60.95 (11.93) | 60.61 (12.16) | 59.31 (13.45) | 65.28 (10.06) | 64.90 (11.06) | 58.26 (15.73) | 61.26 (12.98) |
| gender = male (%) | 304 (73.8) | 12 (1.1) | 386 (73.1) | 346 (64.4) | 255 (67.6) | 242 (46.4) | 102 (55.1) | 290 (61.7) | 1937 (46.9) |
| race (%) | | | | | | | | | |
| american indian or alaska native | 0 (0.0) | 1 (0.1) | 2 (0.4) | 0 (0.0) | 2 (0.5) | 1 (0.2) | 0 (0.0) | 0 (0.0) | 6 (0.1) |
| asian | 44 (10.7) | 61 (5.6) | 11 (2.1) | 8 (1.5) | 161 (42.7) | 8 (1.5) | 11 (5.9) | 12 (2.6) | 316 (7.7) |
| black or african american | 23 (5.6) | 183 (16.7) | 48 (9.1) | 56 (10.4) | 17 (4.5) | 53 (10.2) | 7 (3.8) | 1 (0.2) | 388 (9.4) |
| not reported | 18 (4.4) | 95 (8.7) | 15 (2.8) | 7 (1.3) | 10 (2.7) | 67 (12.8) | 5 (2.7) | 10 (2.1) | 227 (5.5) |
| white | 327 (79.4) | 757 (69.0) | 452 (85.6) | 466 (86.8) | 187 (49.6) | 393 (75.3) | 162 (87.6) | 447 (95.1) | 3191 (77.3) |
| ethnicity (%) | | | | | | | | | |
| hispanic or latino | 9 (2.2) | 39 (3.6) | 26 (4.9) | 26 (4.8) | 18 (4.8) | 7 (1.3) | 5 (2.7) | 11 (2.3) | 141 (3.4) |
| not hispanic or latino | 371 (90.0) | 884 (80.6) | 465 (88.1) | 359 (66.9) | 340 (90.2) | 389 (74.5) | 137 (74.1) | 446 (94.9) | 3391 (82.1) |
| not reported | 32 (7.8) | 174 (15.9) | 37 (7.0) | 152 (28.3) | 19 (5.0) | 126 (24.1) | 43 (23.2) | 13 (2.8) | 596 (14.4) |
| ajcc_pathologic_stage (%) | | | | | | | | | |
| Not Reported | 0 (0.0) | 0 (0.0) | 0 (0.0) | 0 (0.0) | 0 (0.0) | 0 (0.0) | 0 (0.0) | 14 (3.2) | 14 (0.4) |
| Stage 0 | 0 (0.0) | 0 (0.0) | 0 (0.0) | 0 (0.0) | 0 (0.0) | 0 (0.0) | 0 (0.0) | 7 (1.6) | 7 (0.2) |
| Stage I | 2 (0.5) | 183 (16.9) | 27 (6.0) | 269 (50.4) | 175 (49.6) | 279 (54.3) | 21 (11.5) | 77 (17.8) | 1033 (26.1) |
| Stage II | 131 (32.0) | 621 (57.2) | 74 (16.3) | 57 (10.7) | 87 (24.6) | 124 (24.1) | 152 (83.5) | 140 (32.4) | 1386 (35.0) |
| Stage III | 141 (34.4) | 249 (22.9) | 82 (18.1) | 125 (23.4) | 86 (24.4) | 85 (16.5) | 4 (2.2) | 171 (39.6) | 943 (23.8) |
| Stage IV | 136 (33.2) | 20 (1.8) | 270 (59.6) | 83 (15.5) | 5 (1.4) | 26 (5.1) | 5 (2.7) | 23 (5.3) | 568 (14.3) |
| Stage X | 0 (0.0) | 13 (1.2) | 0 (0.0) | 0 (0.0) | 0 (0.0) | 0 (0.0) | 0 (0.0) | 0 (0.0) | 13 (0.3) |
| days_to_death (mean (SD)) | 550.82 (533.62) | 1584.62 (1312.07) | 738.26 (923.07) | 926.72 (741.28) | 669.14 (695.04) | 790.41 (700.86) | 459.27 (362.68) | 1782.98 (1888.91) | 980.13 (1159.26) |
| ajcc_pathologic_t (%) | | | | | | | | | |
| T0 | 1 (0.3) | 0 (0.0) | 1 (0.2) | 0 (0.0) | 0 (0.0) | 0 (0.0) | 0 (0.0) | 23 (5.2) | 25 (0.6) |
| T1 | 3 (0.8) | 281 (25.6) | 49 (9.7) | 275 (51.2) | 185 (49.3) | 172 (33.0) | 7 (3.8) | 42 (9.5) | 1014 (25.1) |
| T2 | 120 (31.6) | 635 (57.9) | 140 (27.7) | 69 (12.8) | 95 (25.3) | 281 (53.8) | 24 (13.0) | 78 (17.7) | 1442 (35.7) |
| T3 | 196 (51.6) | 138 (12.6) | 101 (20.0) | 182 (33.9) | 81 (21.6) | 47 (9.0) | 148 (80.4) | 90 (20.4) | 983 (24.3) |
| T4 | 59 (15.5) | 40 (3.6) | 175 (34.7) | 11 (2.0) | 13 (3.5) | 19 (3.6) | 4 (2.2) | 153 (34.7) | 474 (11.7) |

| | | | | | | | | | |
|---|---|---|---|---|---|---|---|---|---|
| Tis | 0 (0.0) | 0 (0.0) | 0 (0.0) | 0 (0.0) | 0 (0.0) | 0 (0.0) | 0 (0.0) | 8 (1.8) | 8 (0.2) |
| TX | 1 (0.3) | 3 (0.3) | 39 (7.7) | 0 (0.0) | 1 (0.3) | 3 (0.6) | 1 (0.5) | 47 (10.7) | 95 (2.4) |
| ajcc_pathologic_n (%) | | | | | | | | | |
| N0 | 239 (58.9) | 516 (47.0) | 180 (35.8) | 240 (44.7) | 257 (68.4) | 335 (64.3) | 50 (27.2) | 235 (52.3) | 2052 (50.4) |
| N1 | 47 (11.6) | 364 (33.2) | 68 (13.5) | 17 (3.2) | 4 (1.1) | 98 (18.8) | 130 (70.7) | 74 (16.5) | 802 (19.7) |
| N2 | 76 (18.7) | 120 (10.9) | 172 (34.2) | 0 (0.0) | 0 (0.0) | 75 (14.4) | 0 (0.0) | 49 (10.9) | 492 (12.1) |
| N3 | 8 (2.0) | 77 (7.0) | 8 (1.6) | 0 (0.0) | 0 (0.0) | 2 (0.4) | 0 (0.0) | 55 (12.2) | 150 (3.7) |
| NX | 36 (8.9) | 20 (1.8) | 75 (14.9) | 280 (52.1) | 115 (30.6) | 11 (2.1) | 4 (2.2) | 36 (8.0) | 577 (14.2) |
| ajcc_pathologic_m (%) | | | | | | | | | |
| M0 | 196 (47.9) | 912 (83.1) | 191 (74.3) | 426 (79.6) | 272 (72.1) | 353 (68.1) | 85 (45.9) | 418 (94.6) | 2853 (74.7) |
| M1 | 11 (2.7) | 22 (2.0) | 1 (0.4) | 79 (14.8) | 4 (1.1) | 25 (4.8) | 5 (2.7) | 24 (5.4) | 171 (4.5) |
| MX | 202 (49.4) | 163 (14.9) | 65 (25.3) | 30 (5.6) | 101 (26.8) | 140 (27.0) | 95 (51.4) | 0 (0.0) | 796 (20.8) |

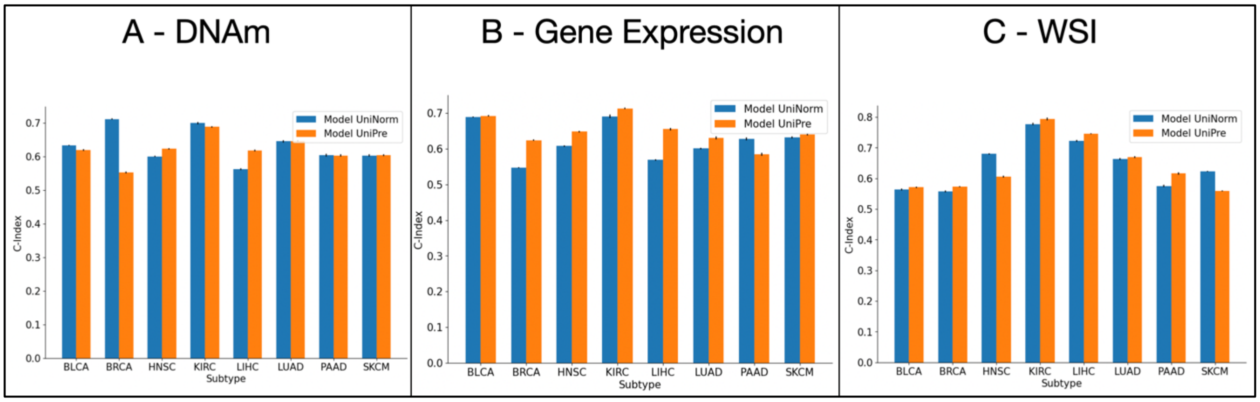


S2**:** Bar graph comparison of unimodal pretraining strategies performances by modality

S3: C-indices of WSI GCNs pretrained using UniPre-WSI with DNAm prediction as the crossmodal prediction target.

| **Subtype** | **C-Index** |
|---|---|
| BLCA | 0.551 ± 0.004 |
| BRCA | 0.600 ± 0.003 |
| HNSC | 0.601 ± 0.002 |
| KIRC | 0.770 ± 0.002 |
| LIHC | 0.741 ± 0.003 |
| LUAD | 0.676 ± 0.003 |
| PAAD | 0.453 ± 0.002 |
| SKCM | 0.562 ± 0.001 |

S4**:** Unimodal boostrapped C-indices with 95% confidence intervals for clinical models, both with and without stage

| **Subtype** | **RF- No Stage** | **RF-Stage** | **CoxPH- No Stage** | **CoxPH-Stage** |
|---|---|---|---|---|
| BLCA | 0.585 ± 0.009 | **0.699 ± 0.006** | 0.590 ± 0.009 | 0.637 ± 0.005 |
| BRCA | 0.511 ± 0.010 | 0.589 ± 0.010 | 0.504 ± 0.011 | **0.590 ± 0.009** |

| | | | | |
|---|---|---|---|---|
| HNSC | 0.542 ± 0.006 | 0.606 ± 0.007 | 0.547 ± 0.007 | **0.609 ± 0.006** |
| KIRC | 0.576 ± 0.011 | 0.784 ± 0.008 | 0.585 ± 0.013 | **0.787 ± 0.008** |
| LIHC | 0.525 ± 0.008 | 0.663 ± 0.008 | 0.531 ± 0.008 | **0.660 ± 0.008** |
| LUAD | 0.465 ± 0.012 | 0.621 ± 0.012 | 0.466 ± 0.012 | **0.622 ± 0.011** |
| PAAD | **0.516 ± 0.007** | 0.497 ± 0.008 | 0.512 ± 0.008 | 0.489 ± 0.008 |
| SKCM | 0.464 ± 0.007 | 0.562 ± 0.012 | 0.466 ± 0.007 | **0.573 ± 0.012** |

S5: 95% Confidence Interval UniTransfer C-indices reported per subtype per method; note that finetuning to each subtype was not performed at the unimodal level in this experiment

| Subtype | UniTransfer-Omics (Gene Expression) | UniTransfer-Omics (DNAM) | UniTransfer-WSI |
|---|---|---|---|
| **BLCA** | 0.674 ± 0.002 | 0.660 ± 0.002 | 0.582 ± 0.003 |
| **BRCA** | 0.742 ± 0.003 | 0.623 ± 0.002 | 0.752 ± 0.002 |
| **HNSC** | 0.626 ± 0.002 | 0.554 ± 0.002 | 0.445 ± 0.002 |
| **KIRC** | 0.552 ± 0.003 | 0.324 ± 0.002 | 0.719 ± 0.003 |
| **LIHC** | 0.582 ± 0.004 | 0.573 ± 0.003 | 0.626 ± 0.004 |
| **LUAD** | 0.610 ± 0.004 | 0.637 ± 0.002 | 0.657 ± 0.003 |
| **PAAD** | 0.590 ± 0.004 | 0.542 ± 0.005 | 0.505 ± 0.004 |
| **SKCM** | 0.577 ± 0.002 | 0.545 ± 0.003 | 0.434 ± 0.002 |

S6: Significance of incorporation of deep learning hazards improving prognostic capability, in addition to using solely clinical covariates and solely pTNM staging, using CoxPH models

| Subtype | H1: Hazard+Stage>Hazard | H2: Hazard+Stage>Stage |
|---|---|---|
| BLCA | 0.057 | 0.120 |
| BRCA | 0.187 | 0.247 |
| HNSC | 0.071 | 0.085 |
| KIRC | 0.034 | 0.111 |
| LIHC | 0.148 | 0.119 |
| LUAD | 0.132 | 0.123 |
| PAAD | 0.168 | 0.319 |
| SKCM | 0.030 | 0.099 |

S7: Most significant pathways identified by Integrated Gradients on gene expression encoders of top performing multimodal models

| BLCA | BRCA | HNSC | KIRC | LIHC | LUAD | PAAD | SKCM |
|---|---|---|---|---|---|---|---|
| positive regulation of mitotic nuclear division (GO:0045840) | regulation of cellular localization (GO:0060341) | defense response to Gram-negative bacterium (GO:0050829) | positive regulation of actin nucleation (GO:0051127) | positive regulation of vasoconstriction (GO:0045907) | killing by host of symbiont cells (GO:0051873) | killing by host of symbiont cells (GO:0051873) | skeletal system development (GO:0001501) |
| regulation of smooth muscle cell proliferation (GO:0048660) | supramolecular fiber organization (GO:0097435) | mesoderm formation (GO:0001707) | regulation of mitotic nuclear division (GO:0007088) | negative regulation of Wnt signaling pathway (GO:0030178) | antibacterial humoral response (GO:0019731) | regulation of cell-matrix adhesion (GO:0001952) | killing by host of symbiont cells (GO:0051873) |

| | | | | | | | |
|---|---|---|---|---|---|---|---|
| kidney development (GO:0001822) | mammary gland development (GO:0030879) | intermediate filament bundle assembly (GO:0045110) | translational elongation (GO:0006414) | negative regulation of production of molecular mediator of immune response (GO:0002701) | antimicrobial humoral immune response mediated by antimicrobial peptide (GO:0061844) | receptor clustering (GO:0043113) | positive regulation of non-canonical Wnt signaling pathway (GO:2000052) |
| regulation of type 2 immune response (GO:0002828) | Fc receptor mediated stimulatory signaling pathway (GO:0002431) | regulation of leukocyte activation (GO:0002694) | receptor catabolic process (GO:0032801) | heme biosynthetic process (GO:0006783) | organic hydroxy compound transport (GO:0015850) | regulation of pri-miRNA transcription by RNA polymerase II (GO:1902893) | epidermis development (GO:0008544) |
| development of primary male sexual characteristics (GO:0046546) | regulation of macrophage activation (GO:0043030) | chemical synaptic transmission (GO:0007268) | negative regulation of actin filament polymerization (GO:0030837) | cellular response to starvation (GO:0009267) | positive regulation of cellular process (GO:0048522) | muscle filament sliding (GO:0030049) | positive regulation of cell differentiation (GO:0045597) |
| organic hydroxy compound transport (GO:0015850) | regulation of heterotypic cell-cell adhesion (GO:0034114) | response to interferon-gamma (GO:0034341) | positive regulation of NIK/NF-kappaB signaling (GO:1901224) | iron ion homeostasis (GO:0055072) | skeletal system development (GO:0001501) | hydrogen peroxide biosynthetic process (GO:0050665) | positive regulation of cellular process (GO:0048522) |
| terpenoid metabolic process (GO:0006721) | regulation of cysteine-type endopeptidase activity (GO:2000116) | hemidesmosome assembly (GO:0031581) | positive regulation of neuron death (GO:1901216) | very-low-density lipoprotein particle remodeling (GO:0034372) | cellular metal ion homeostasis (GO:0006875) | Fc receptor signaling pathway (GO:0038093) | adenylate cyclase-modulating G protein-coupled receptor signaling pathway (GO:0007188) |
| sensory perception of sound (GO:0007605) | alpha-linolenic acid metabolic process (GO:0036109) | establishment of skin barrier (GO:0061436) | regulation of phagocytosis, engulfment (GO:0060099) | zymogen activation (GO:0031638) | cellular response to fatty acid (GO:0071398) | positive regulation of cell population proliferation (GO:0008284) | antimicrobial humoral immune response mediated by antimicrobial peptide (GO:0061844) |
| organic acid transport (GO:0015849) | negative regulation of cell motility (GO:2000146) | intermediate filament organization (GO:0045109) | autophagy of peroxisome (GO:0030242) | protein processing (GO:0016485) | C21-steroid hormone metabolic process (GO:0008207) | defense response to Gram-positive bacterium (GO:0050830) | regulation of focal adhesion assembly (GO:0051893) |
| organic heteropentacyclic compound metabolic | innate immune response in | regulation of water loss via skin (GO:0033561) | maintenance of blood-brain barrier | epithelial cell differentiation (GO:0030855) | positive regulation of granulocyte chemotaxis | positive regulation of coagulation | skin development (GO:0043588) |

| | | | | |
|---|---|---|---|---|
| process (GO:1901376) | mucosa (GO:0002227) | (GO:0035633 ) | (GO:0071624 ) | (GO:0050820 ) |

S8: Most significant genes identified by Integrated Gradients on DNA methylation encoders of top performing multimodal models

| **BLCA** | **BRCA** | **HNSC** | **KIRC** | **LIHC** | **LUAD** | **PAAD** | **SKCM** |
|---|---|---|---|---|---|---|---|
| GSTA3 | IMPDH2 | CLPS | PDE11A | EEF1D | DAZAP1 | SS18L1 | IGFL2 |
| RAX | TOR2A | BCAP31 | DOK7 | JRK | LOC284788 | VDAC1 | RP2 |
| LOC100190940 | FPR2 | LOC340074 | CDH15 | OTOP2 | RAC2 | HCFC1R1 | HRCT1 |
| ZNF214 | SSBP3 | MDH1B | CLDN16 | EP400 | ARHGEF11 | VGLL1 | KDM2B |
| TEPP | CLEC17A | CNTFR | DAXX | VASN | LILRP2 | UXT | PCDH15 |
| PLOD2 | PPP1CC | ABL2 | KNDC1 | KIAA0114 | ZCRB1 | LILRP2 | GOLGA8A |
| GNAS | GNG7 | TMEM102 | FRMD4A | IDS | LOC729156 | MAGOH | PI4K2A |
| EFCAB6 | INPP5D | NCAM1 | VANGL2 | NR2E1 | ICOS | SNORD62B | ZCRB1 |
| GATA2 | SCRN3 | THRA | HOXC6 | ZFP41 | MAGOH | ICOS | MAGOH |
| ZNF354C | CCNA2 | PDK2 | PRDM16 | NSUN4 | CCNA2 | CCNA2 | SNORD62B |

S9: Enrichr pathway analysis of reported significant genes from DNAm data, using BioPlanet pathways

| **Subtype** | **Pathway** | **Adj. P-Value** |
|---|---|---|
| **BLCA** | Rapid glucocorticoid receptor pathway | 0.004492 |
| | Attenuation of GPCR signaling | 0.005488 |
| | Beta-arrestins in GPCR desensitization | 0.005985 |
| | LPA4-mediated signaling events | 0.007973 |
| | Beta-arrestin-dependent recruitment of Src kinases in GPCR signaling | 0.008469 |
| | Ion channel function in vascular endothelium | 0.008469 |
| | Corticosteroids and cardioprotection | 0.008966 |
| | Serotonin receptor 4/6/7 and NR3C signaling | 0.008966 |
| | PKA activation in glucagon signaling | 0.008966 |
| | GATA3-mediated activation of Th2 cytokine expression | 0.009957 |
| **BRCA** | G2 phase pathway | 0.002498 |
| | Formyl peptide interaction with formyl peptide receptors | 0.003994 |
| | Purine ribonucleoside monophosphate biosynthesis | 0.005488 |
| | Platelet endothelial cell adhesion molecule 1 (PECAM1) interactions | 0.005985 |
| | Hormone-sensitive lipase (HSL)-mediated triacylglycerol hydrolysis | 0.005985 |
| | PI3K class IB pathway | 0.006482 |
| | Cyclin A/B1-associated events during G2/M transition | 0.007476 |
| | Presynaptic function of kainate receptors | 0.01045 |
| | T cell receptor downstream signaling | 0.01095 |
| | Signaling events mediated by PRL | 0.01144 |
| **HNSC** | Vitamin A uptake in enterocytes | 0.003495 |
| | Abl role in Robo-Slit signaling | 0.004492 |
| | Pyruvate dehydrogenase (PDH) complex regulation | 0.005985 |
| | MAP kinase inactivation of SMRT corepressor | 0.006979 |
| | Apoptotic cleavage of cellular proteins | 0.008469 |
| | Synaptic proteins at the synaptic junction | 0.008469 |
| | PTEN-dependent cell cycle arrest and apoptosis | 0.009462 |
| | Skeletal muscle hypertrophy is regulated via AKT/mTOR pathway | 0.009957 |
| | Y branching of actin filaments | 0.01045 |
| | RXR/VDR pathway | 0.01293 |
| **KIRC** | Stress induction of HSP regulation | 0.007476 |
| | Transcriptional activity regulation by PML | 0.008966 |
| | Cell junction organization | 0.0007674 |
| | Nitric oxide stimulation of guanylate cyclase | 0.01392 |
| | Adherens junction actin cytoskeletal organization | 0.01441 |
| | CDO in myogenesis | 0.01441 |
| | Tight junction interactions | 0.01490 |

| | | |
|---|---|---|
| | Cell-cell communication | 0.001796 |
| | Cell adhesion molecules (CAMs) | 0.001907 |
| | FAS pathway and stress induction of heat shock protein regulation | 0.01933 |
| **LIHC** | Chondroitin sulfate/dermatan sulfate degradation | 0.006482 |
| | Glycosaminoglycan degradation | 0.009462 |
| | Heparan sulfate/heparin glycosaminoglycan (HS-GAG) degradation | 0.009957 |
| | Nuclear receptors | 0.01884 |
| | Chondroitin sulfate/dermatan sulfate metabolism | 0.02375 |
| | Translation factors | 0.02473 |
| | Nuclear receptor transcription pathway | 0.02521 |
| | Heparan sulfate/heparin glycosaminoglycan (HS-GAG) metabolism | 0.02570 |
| | Glycosaminoglycan metabolism | 0.05367 |
| | Lysosome | 0.05889 |
| **LUAD** | G2 phase pathway | 0.002498 |
| | Sema4D in semaphorin signaling | 0.00009070 |
| | Cyclin A/B1-associated events during G2/M transition | 0.007476 |
| | Semaphorin interactions | 0.0004745 |
| | STAT3 pathway | 0.009462 |
| | T cell activation co-stimulatory signal | 0.01045 |
| | G alpha (12/13) signaling events | 0.0006453 |
| | Signaling events mediated by PRL | 0.01144 |
| | Rho cell motility signaling pathway | 0.01144 |
| | G0 and early G1 pathway | 0.01243 |
| **PAAD** | G2 phase pathway | 0.002498 |
| | Cyclin A/B1-associated events during G2/M transition | 0.007476 |
| | E2F transcription factor network | 0.0005962 |
| | T cell activation co-stimulatory signal | 0.01045 |
| | Signaling events mediated by PRL | 0.01144 |
| | G0 and early G1 pathway | 0.01243 |
| | Interleukin-2/STAT5 pathway | 0.01490 |
| | Primary immunodeficiency | 0.01737 |
| | FRA pathway | 0.01835 |
| | FOXM1 transcription factor network | 0.02032 |
| **SKCM** | PIP biosynthesis at the early endosome membrane | 0.006482 |
| | PIP biosynthesis at the Golgi membrane | 0.008469 |
| | PIP biosynthesis at the plasma membrane | 0.01638 |
| | Cleavage of growing transcript in the termination region | 0.02130 |
| | Phosphatidylinositol metabolism | 0.02473 |
| | Transport of mature transcript to cytoplasm | 0.02717 |
| | Inositol phosphate metabolism | 0.02814 |
| | Messenger RNA splicing: major pathway | 0.03349 |
| | Phosphatidylinositol signaling system | 0.03833 |
| | RNA polymerase II transcription | 0.04938 |

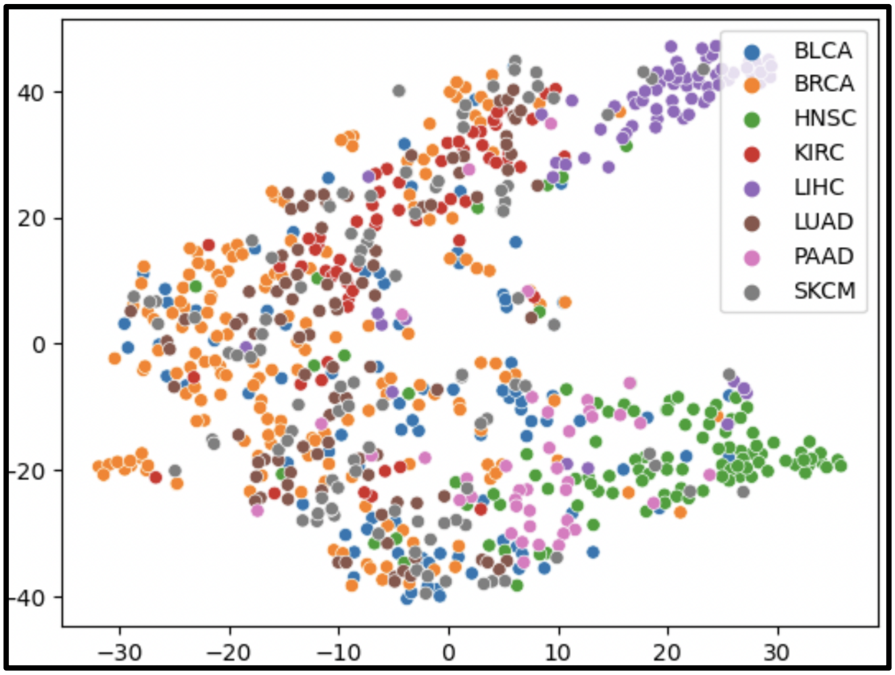


S10: Visualized t-SNE embeddings of fused features extracted from MultiTrans prior to subtype finetuning

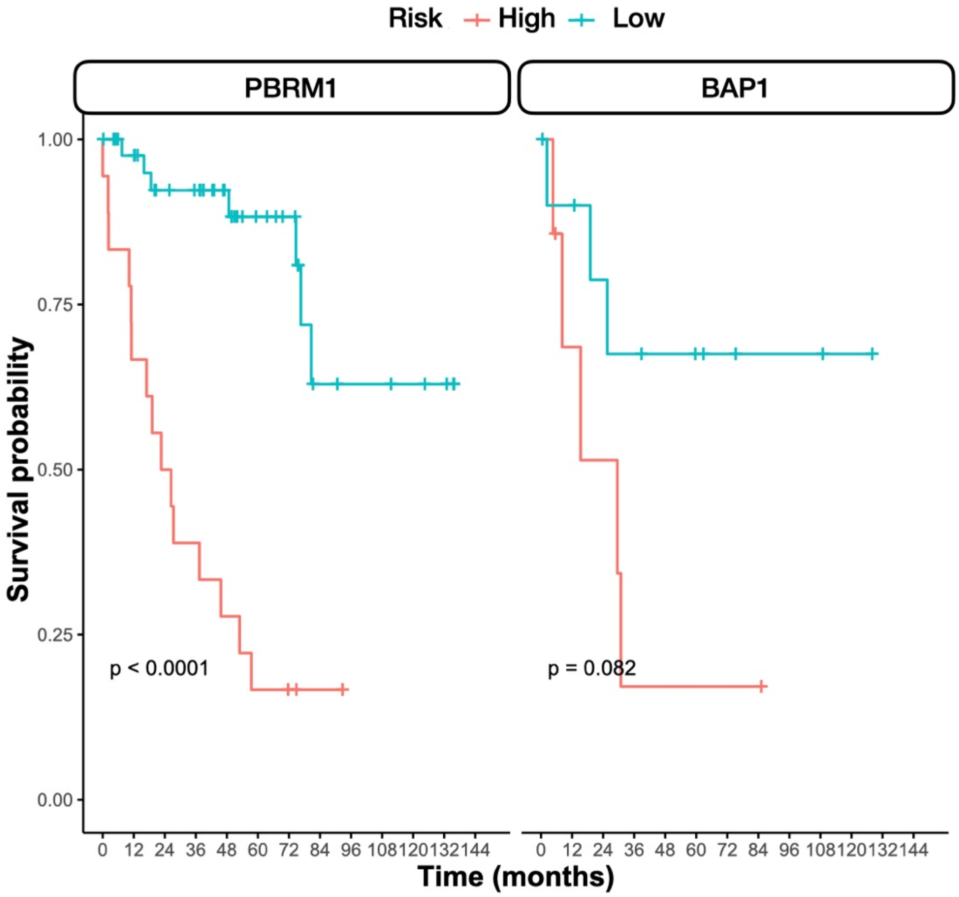

S11: Kaplan Meier curves for predicted risk, stratified by patients with the following KIRC molecular subtypes: A) PBRM1 mutation, B) BAP1 mutation

S12**:** Relationship between the ability of the multimodal models to localize TILs and predicted risk

| **Subtype** | **Odds Ratio** | **P-Value** |
|---|---|---|
| BLCA | 0.44 | 0.0029 |
| BRCA | 0.56 | 0.1208 |
| LUAD | 1.76 | 0.0132 |
| PAAD | 0.33 | 0.0087 |
| SKCM | 2.11 | 0.0187 |

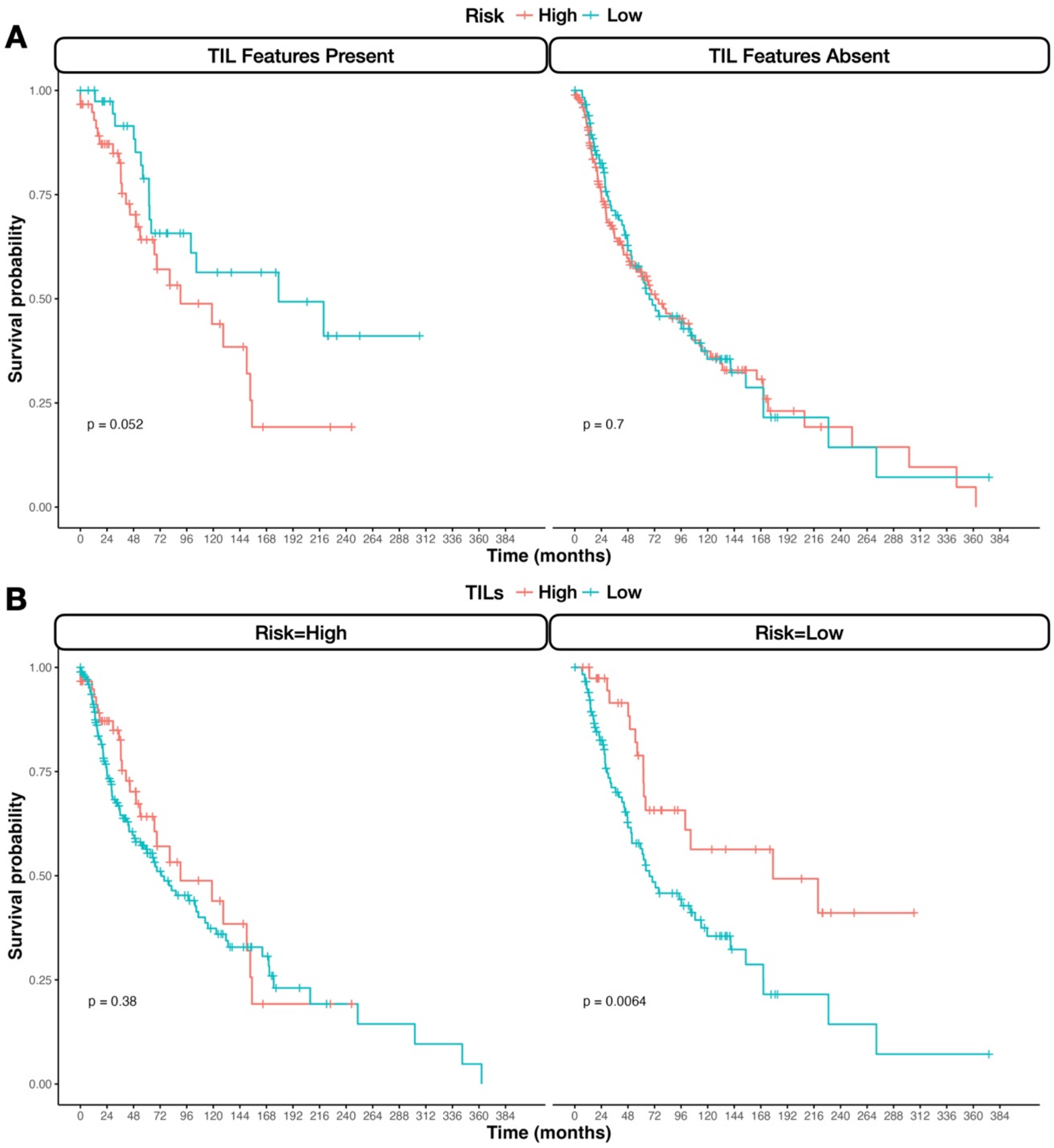

S13: Kaplan Meier survival curves depicting interaction between TIL identification by multimodal model and predicted risk: A) Indicators of high or low predicted risk, stratified by model localization of TILs, B) Model localization of TILs, stratified by indicators of high or low predicted risk

S14: Integrated gradients scores for significant genes derived from top-performing multimodal model DNAm networks. **See excel file, S14_DNAm_IG.xlsx**

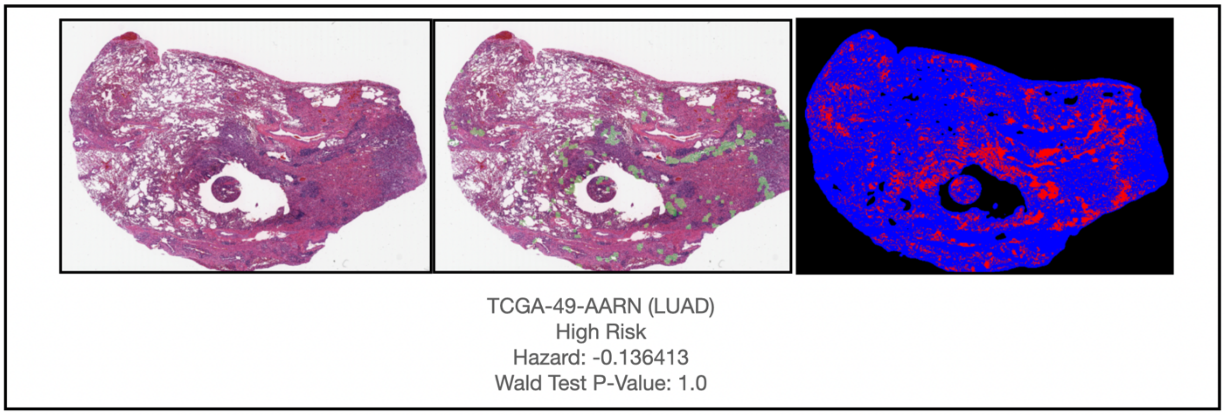


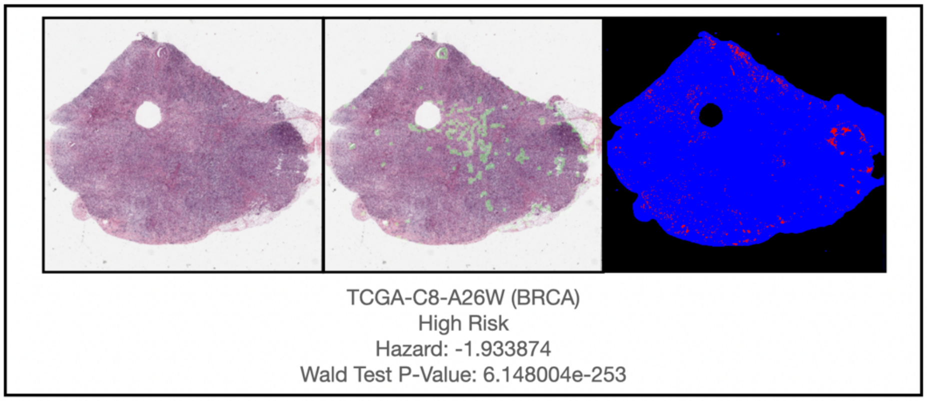


S15: Additional interpretation of WSI GCNs from top performing multimodal models

S16: Size of train, test, and validation splits, per subtype

| Subtype | Train | Validation | Test |
|---|---|---|---|
| **BLCA** | 291 | 45 | 46 |
| **BRCA** | 581 | 90 | 90 |
| **HNSC** | 338 | 52 | 53 |
| **KIRC** | 226 | 35 | 36 |
| **LIHC** | 274 | 42 | 43 |
| **LUAD** | 318 | 49 | 50 |
| **PAAD** | 134 | 21 | 21 |
| **SKCM** | 330 | 51 | 51 |

S17: Selected hyperparameters for trained models

| Model | Learning Rate | Epochs | Batch Size | Weight Decay |
|---|---|---|---|---|
| UniNorm-Omics VAE | 0.008 | 500 | 32 | 1e-4 |

| | | | | |
|---|---|---|---|---|
| UniNorm-Omics Survival Model | 0.0002 | 40 | 32 | 1e-4 |
| UniNorm-WSI | 0.0001 | 40 | 4*4 | 1e-4 |
| UniPre-Omics | 0.0002 | 40 | 32 | 1e-4 |
| UniPre-WSI | 0.0001 | 40 | 4*4 | 1e-4 |
| MultiNorm | 0.0001 | 40 | 3*8 | 1e-4 |
| MultiPre | 0.0001 | 40 | 3*8 | 1e-4 |
| UniTransfer-Omics | 0.0002 | 40 | 32 | 1e-4 |
| UniTransfer-WSI | 0.0001 | 40 | 4*4 | 1e-4 |
| MultiTransfer | 0.0001 | 10 | 3*8 | 1e-4 |
| MultiTransfer (finetuning) | 0.0001 | 40 | 3*8 | 1e-4 |
